# CRMP2 mediates Sema3F-dependent axon pruning and dendritic spine remodeling

**DOI:** 10.1101/719617

**Authors:** Jakub Ziak, Romana Weissova, Kateřina Jeřábková, Martina Janikova, Roy Maimon, Tomas Petrasek, Barbora Pukajova, Mengzhe Wang, Monika S. Brill, Marie Kleisnerova, Petr Kasparek, Xunlei Zhou, Gonzalo Alvarez-Bolado, Radislav Sedlacek, Thomas Misgeld, Ales Stuchlik, Eran Perlson, Martin Balastik

**Author notes:** All correspondence should be addressed to Martin Balastik. These authors contributed equally (Romana Weissova, Kateřina Jeřábková).

## Abstract

Regulation of axon guidance and pruning of inappropriate synapses by class 3 semaphorins is key to development of neural circuits. Collapsin response mediator protein 2 (CRMP2) has been shown to regulate axon guidance by mediating Semaphorin 3A (Sema3A) signaling and its dysfunction has been linked to schizophrenia, however, nothing is known about its role in the synapse pruning. Here, using newly generated *crmp2^−/−^* mice we demonstrate that while CRMP2 has only a moderate effect on Sema3A-dependent axon guidance *in vivo*, it is essential for Sema3F-dependent axon pruning and dendritic spine remodeling. We first demonstrate that CRMP2 deficiency interferes with Sema3A signaling in compartmentalized neuron cultures and leads to a mild defect in axon guidance in peripheral nerves and corpus callosum. Strikingly, we show that *crmp2^−/−^* mice display more prominent defects in dendritic spine pruning and stereotyped axon pruning in hippocampus and visual cortex consistent with impaired Sema3F signaling and with autism spectrum disorder (ASD)-rather than schizophrenia-like phenotype. Indeed, we demonstrate that CRMP2 mediates Sema3F-induced axon retraction in primary neurons and that *crmp2^−/−^* mice display early postnatal behavioral changes linked to ASD. In conclusion, we demonstrate that CRMP2 is an essential mediator of Sema3F-dependent synapse pruning and its dysfunction in early postnatal stages shares histological and behavioral features of ASD.

## INTRODUCTION

The pattern of axonal connections is established during pre- and postnatal development by a cascade of multiple events. In embryogenesis, axonal growth cones are guided to their targets and multiple axon branches are formed. Since both correct and incorrect projections are formed, the embryonic brain connectome is only transient and the inaccurate connections are eliminated (pruned) in the early postnatal development (Riccomagno and Kolodkin, 2015). Defects in development and maturation of brain circuits have been linked to several neurodevelopmental disorders including autism spectrum disorder (ASD), schizophrenia, or epilepsy (Johnston, 2004, Vanderhaeghen and Cheng, 2010, Van Battum et al., 2015).

Generally, two types of pruning are recognized: 1) small-scale axon pruning, regulated by neural activity or trophic support and 2) large-scale stereotyped axon pruning, which is genetically predetermined (Riccomagno and Kolodkin, 2015, Vanderhaeghen and Cheng, 2010). Stereotyped pruning can be further histologically divided into degeneration-like (Yu and Schuldiner, 2014) and retraction-like (Bagri et al., 2003), which has been linked to secreted semaphorins and their co-receptors e.g. Plexin-A4 and -A3 (Bagri et al., 2003, Low et al., 2008). Intracellular mediators that transmit signals from Plexins in axon pruning are not completely understood. One of the key molecule downstream of Semaphorin 3A (Sema3A) signaling that directly interacts with cytoskeleton components is collapsin response mediator protein 2 (CRMP2) (Goshima et al., 1995, Byk et al., 1996, Minturn et al., 1995). In its non-phosphorylated state, CRMP2 binds to tubulin dimers and promotes their polymerization (Fukata et al., 2002). However, upon phosphorylation, it dissociates from microtubules promoting growth cone collapse (Uchida et al., 2005). Two splice variants of Crmp2 have been found that differ at the N terminus – CRMP2A and CRMP2B (Yuasa-Kawada et al., 2003, Balastik et al., 2015). CRMP2B is phosphorylated by cyclin dependent kinase 5 (CDK5) at Ser522, by GSK-3β at Thr509, 514, 518 and by Rho kinase at Thr555 (Yoshimura et al., 2005, Uchida et al., 2005, Arimura et al., 2005). We have recently shown that CRMP2A is phosphorylated at N-terminal Ser27 by Cdk5, which promotes its interaction and stabilization by prolyl isomerase Pin1 (Balastik et al., 2015). Recently, CRMP2 deficiency *in vivo* has been found to lead to changes in dendritic morphology including decreased spine number and dendritic branches in CA1 pyramidal neurons and layer 5 cortical neurons in conditional and full CRMP2 knockout mice (Zhang et al., 2016, Makihara et al., 2016). Behavioral analysis of the conditional CRMP2 knockout mice revealed hyperactivity and social behavior impairment together with prepulse inhibition (PPI) deficit previously linked to schizophrenia (Zhang et al., 2016). In humans, deregulation of CRMP2 has been associated with several neurological and psychiatric disorders (Quach et al., 2015): in addition to schizophrenia (Nakata et al., 2003, Liu et al., 2014, Clark et al., 2006), CRMP2 has been implicated in Autism Spectrum Disease (ASD) (Braunschweig et al., 2013, De Rubeis et al., 2014), (Sfari Gene database). The exact role of CRMP2 in the development of these conditions has so far been elusive. Although schizophrenia and ASD share some behavioral characteristics (e.g. decreased cognitive functions, impaired social skills, repetitive behavior), they differ in the timing of their onset (early childhood for ASD, late adolescence for schizophrenia) (Lord et al., 2018)) and the nature of the underlying neuronal connectivity disorder. Whereas hypoconnectivity (lower number of dendrites, dendritic spines, general decrease of white matter) is often present in schizophrenia, ASD has been associated with local hyperconectivity caused by either increased synapse formation or their incomplete pruning (Bourgeron, 2015). Because CRMP2 is downstream of Semaphorin signaling, which controls axonal pruning (Bagri et al., 2003), and it has been linked to both schizophrenia and ASD, it is one of the prime candidates to regulate this process. However, the *in vivo* analysis of both conditional and full CRMP2 knockout mice has to date mainly focused on the dendritic phenotype and associated behavioral aspects. In addition, the role of CRMP2 in class 3 semaphorin signaling (other than Sema3A), which has been previously linked to defects in pruning and ASD (particularly Sema3F) is so far not known.

In order to characterize the function of CRMP2 in axon growth, guidance and pruning and its role in class 3 semaphorin signaling *in vivo*, we generated CRMP2 full knockout mice (*crmp2^−/−^*). We show that CRMP2 is critical for axon guidance in both central and peripheral nervous systems. In peripheral nervous system deficiency of CRMP2 leads to mild overgrowth and increased branching of ophthalmic branch of trigeminal nerve and other selected peripheral nerves. In the central nervous system, we detected defects in postnatal callosal axon growth and guidance. Both of these systems are regulated by Sema3A, which was previously shown to induce CRMP2 phosphorylation and signaling *in vitro*. Indeed, we confirm that primary motor neurons isolated from *crmp2^−/−^* mice have defects mediating Sema3A signaling.

Importantly, we show that CRMP2 is essential also for synaptic refinement as *crmp2^−/−^* mice demonstrate defective stereotyped pruning of axons arising from hippocampus and visual cortex and inadequate elimination of dendritic spines in dentate gyrus (DG). Pruning in both of these systems is dependent on Sema3F rather than Sema3A and its defect is in accord with ASD-rather than schizophrenia-like phenotype. In agreement with this hypothesis we show that CRMP2 is essential for Sema3F-induced axon retraction in primary hippocampal cultures and that *crmp2^−/−^* mice suffer from ultrasonic vocalizations defect in early postnatal stages linked to ASD and postnatal cognitive deficit.

In summary, we provide evidence that in addition to its role in Sema3A-dependent axon guidance CRMP2 is a key mediator of Sema3F-dependent axon pruning and dendritic spine remodeling. Our data highlight the importance of CRMP2 in neural circuit formation and refinement *in vivo* and demonstrates that its deficiency leads to defects in neural development associated with neurodevelopmental disorders, in particular ASD and schizophrenia.

## RESULTS

### Crmp2 deficiency leads to axonal growth defects in peripheral nerves

We used TALEN (transcription activator-like effector nucleases) mutagenesis to delete the second and third exons of the *Crmp2* locus leading to a knockout of both CRMP2 protein isoforms CRMP2A and CRMP2B (Fig. 1A, B, Fig. S1A, B). The knockout mice were viable and fertile and the size of their brains was similar to the WT littermates (Fig. S1C). In agreement with (Zhang et al., 2016), we observed ventriculomegaly in the homozygous mice. (Fig. S1D, E).

**Figure 1.**
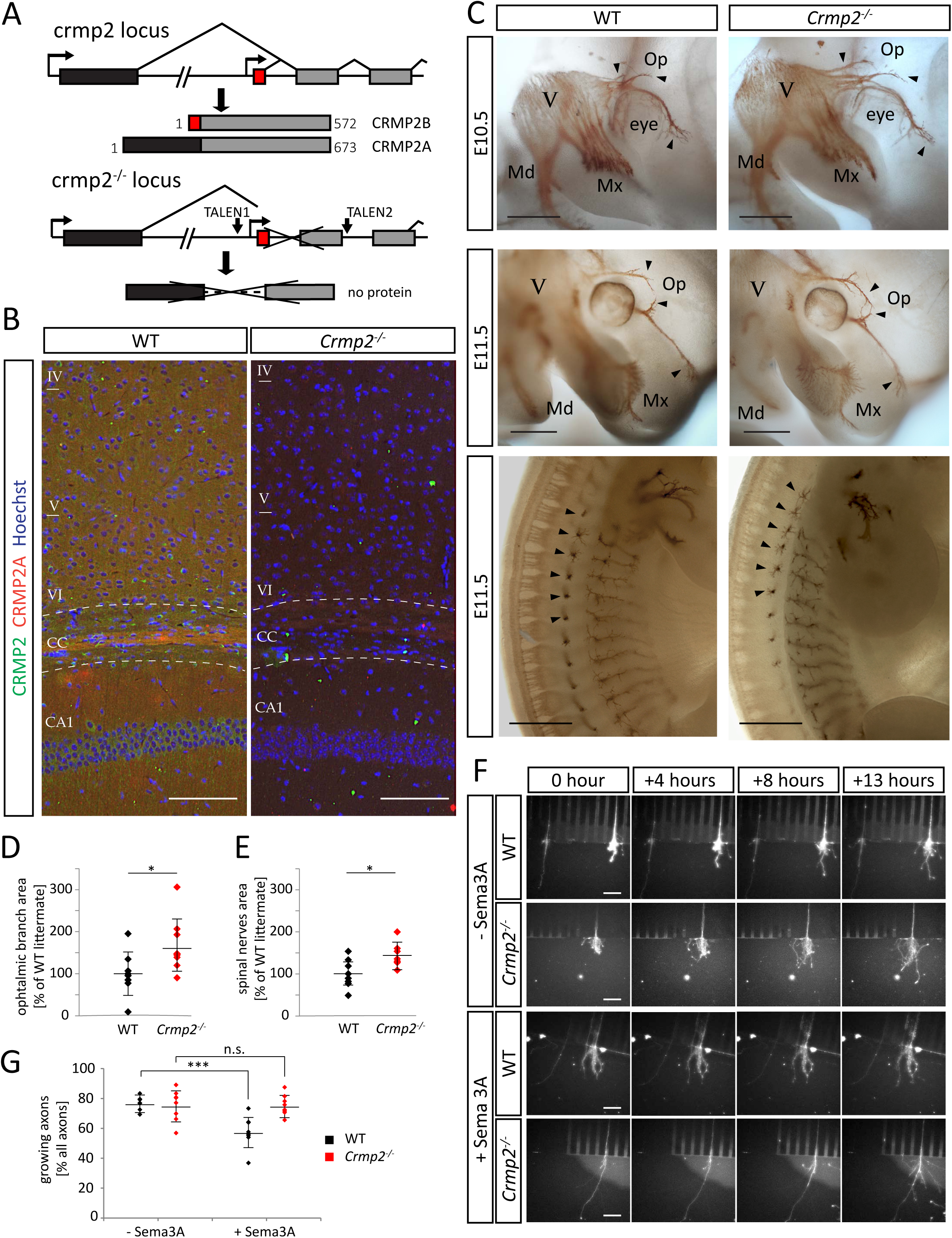
CRMP2 deficiency leads to axonal growth defects in peripheral nerves. (A) Generation of crmp2 knockout mice by TALEN mutagenesis. Alternative exons are highlighted in black and red. (B) CRMP2 expression in adult cortex and corpus callosum. CRMP2B (green) is expressed throughout the cortex and hippocampus (area CA1), CRMP2A (red) is particularly strongly expressed in a subset of callosal axons. Nuclei are counterstained with Hoechst 33342. (C) Increased growth of ophthalmic branch of the trigeminal nerve together with increased defasciculation in E10.5 and E11.5 *crmp2^−/−^* embryos (first and second row, arrowheads). Increased growth and branching of lateral branches of spinal nerves in *crmp2^−/−^* embryos (third row, arrowheads). (D, E) Quantification of areas innervated by ophthalmic branches of trigeminal ganglions and spinal nerves, normalized to WT littermates (each dot depicts one embryo). Area of ophthalmic branch in *crmp2^−/−^* was increased by 66% (p<0.05, n=10, 3 litters). For spinal nerves, total area in knockouts was increased by 45% (p<0.05, WT n=8, *crmp2^−/−^* n=7, 3 litters) (F) Growth of motor neurons from E11.5 – E12.5 WT and *crmp2^−/−^* spinal cord explants in microfluidic chambers. Excerpts from 14 hour live imaging. Upon application of Sema3A (5nM), WT axons tend to stop or slow down their growth unlike *crmp2^−/−^* axons. (G) Quantification of the fraction of growing (>50 μm) axons in one imaging field (WT control 76.8±5.7%, Sema3A: 57.7±10.6% p<0.001, *crmp2^−/−^* control 75.4±10.4%, Sema3A 75.4±6.7%, p=0.99), n=3 experiments per genotype, 7-8 explants per condition). Data are present as means±SD. * p<0.05, *** p<0.001, t-test. Scale bars: 50 μm in B, 500 μm in C, 100 μm in F. See also Supplementary video 1 and Fig. S1. and S2.

CRMP2 has been shown to mediate Sema3A signaling and regulate axon guidance *in vitro* (Goshima et al., 1995). To test whether CRMP2 deficiency leads to altered axon growth also *in vivo*, we first analyzed development of peripheral nerves in E10.5 – E12.5 embryos using whole-mount immunohistochemistry (Fig. S2A) as their development is regulated by Sema3A (Kitsukawa et al., 1997, Taniguchi et al., 1997, Chen et al., 2000, Cheng et al., 2001). We quantified changes in neuron growth by measuring the area innervated by axons from each given nerve and branching by counting of the total number of branches. The growth and branching of ophthalmic branch of the trigeminal nerve was significantly increased in *crmp2^−/−^* mice at E10.5 – E12.5 and similarly increased was the sprouting of the lateral branches of the spinal nerves (Axon projections from dorsal root ganglions (DRGs) were not changed in *crmp2^−/−^* mice) (Fig. 1C – E, Fig. S2B, C). Our data indicate that the growth-promoting function of CRMP2 is redundant *in vivo* at least in the tested neurons, but that its function as a mediator of repulsive axon guidance signals is unique and cannot be fully rescued by other proteins. Since growth of both the ophthalmic branch as well as the lateral branches of the spinal nerves are negatively regulated by Sema3A (Taniguchi et al., 1997), our data support the function of CRMP2 as a mediator of Sema3A signaling.

To directly demonstrate that CRMP2 deficiency interferes with Sema3A signaling in peripheral neurons isolated from *crmp2^−/−^* mice, we took an advantage of microfluidic chamber approach for extraaxonal environment manipulation. In this chamber, cell bodies and axons are separated in proximal or distal compartment, respectively (Maimon et al., 2018). We prepared spinal cord explants from WT and *crmp2^−/−^* E11.5 – E12.5 embryos and cultured them for 4 – 5 days until motor neuron axons crossed to the distal compartment (Fig. S2D). Then, we stained axons with Alexa 647-conjugated choleratoxin subunit B and analyzed their growth by live imaging. Without Sema3A addition the growth of WT and *crmp2^−/−^* explants and their axons was comparable indicating again that the axon growth-promoting function of CRMP2 is redundant, at least in the tested neurons. When we added Sema3A (5nM) to the distal (axonal) compartment, we observed decrease in the number of growing WT axons, as expected (p<0.001, Fig. 1F, G video 1). However, in *crmp2^−/−^* explants, Sema3A had no significant effect on the number of growing axons (p=0,99, Fig. 2F, G, video 1). Thus, using both *in vitro* and *in vivo* analysis we demonstrate that CRMP2 mediates Sema3A signaling in peripheral nerves in early development, which is in accord with the previously published *in vitro* data (Goshima et al., 1995, Brown et al., 2004).

**Figure 2.**
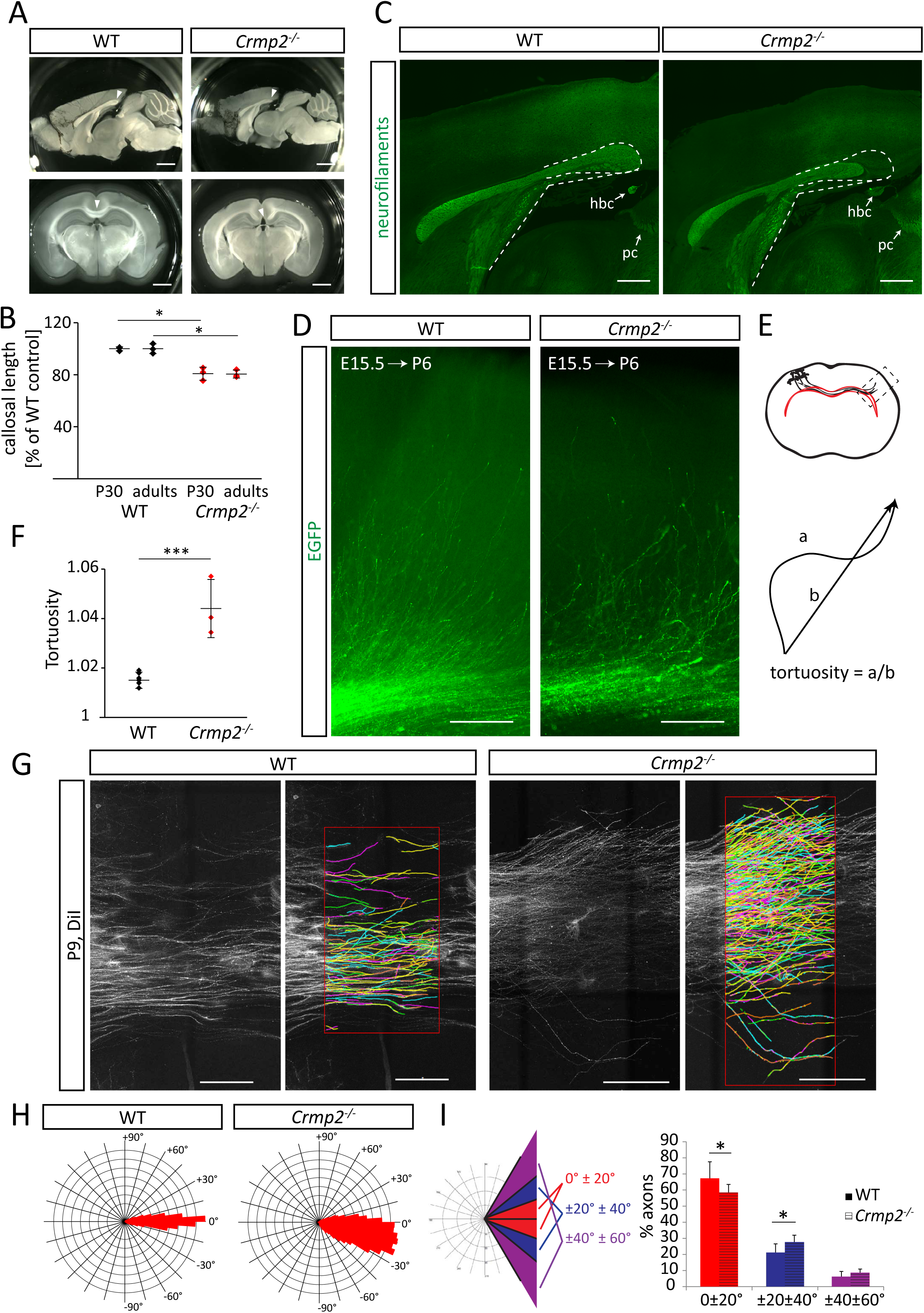
CRMP2 regulates postnatal development of corpus callosum. (A) CRMP2 deficiency leads to callosal hypoplasia. Shortening of caudal corpus callosum (arrowheads) is apparent in both sagittal (first row) and coronal sections (second row) of adult brains. (B) Quantification of callosal length in 30-days old (P30, n=3, knockout is 80.8±5% of WT, p=0.003) and adult mice (n=3, 80.5±3% of WT, p=0.002), means±SD are shown. (C) Labeling of adult corpus callosum with anti-neurofilaments antibody in sagittal sections. Outline depicts missing posterior part of the tract in *crmp2^−/−^* mice. Caudal part of the corpus callosum in WTs is located dorsally above the habenular commissure (hbc), while in *crmp2^−/−^* mice callosum terminates rostrally before reaching the hbc. pc – posterior commissure. (D) The growth of GFP-labeled layer 2/3 cortical axons into the contralateral cortex at P6 (electroporated at E15.5, WT n=6, *crmp2^−/−^* n=3). Note the disorganized growth of *crmp2^−/−^*axons. (E) Schematic drawing of the axon growth of the cortical layer 2/3 neuron into the contralateral cortex (rectangle depicts the area displayed in (D)) and explanation of tortuosity parameter. Tortuosity=1 if a=b. (F) Quantification of tortuosity of axons upon their exit from the callosal tract (WT 1.016±0.003 vs. *crmp2^−/−^* 1.044±0.012, p<0.001), means±SD are shown. (G) Maximal projections of DiI-labeled callosal axons from P9 oblique sections and their reconstruction in Neurolucida 360 (WT n=6, c*rmp2^−/−^* n=9). (H) Polar histograms of callosal axons reconstructed in (G). Note the broader range of axon growth angles in *crmp2^−/−^* mice. (I) Left: schematic representation of polar histogram analysis by clustering the traced axons into 3 groups based on the growth angles. Right: proportion of axons growing in selected clusters (WT, 0°-±20°: 67.3±10.3%; ±20°-±40°: 21.2±5.3%; ±40°-±60°: 6.22±3.2%; *crmp2^−/^*^−^, 58.5±5%, p=0.05, 27.7±4.3%, p=0.02, 8.64±2.25%, p=0.1), means±SD are shown. * p<0.05, ***<0.001, t-test. Scale bars: 1 mm in A, 500 µm in B, 200 μm in D, 200 μm in G. See also Fig. S3.

### CRMP2 regulates anatomy and axon guidance in Corpus callosum

In addition to the peripheral nerves, we analyzed the effect of CRMP2 deficiency on axonal growth also in the central nervous system. We detected anatomical changes in the largest axonal bundle of the brain, the Corpus callosum, in *crmp2^−/−^* mice. The length of Corpus callosum was significantly reduced in juvenile (−19.2%, p<0,05) as well as adult (−19.5%, p<0,05) knockout mice as can be appreciated in both sagittal and coronal sections (Fig. 2A, B). More detailed labeling of the tract with anti-neurofilaments antibody showed that the posterior part of the Corpus callosum (splenium) is markedly hypoplastic in *crmp2^−/−^* mice, ending rostrally to the habenular commissure, while the WT splenium is longer, located just above the habenular commissure (Fig. 2C).

Callosal axons have been shown to be guided by Sema3A (Zhou et al., 2013, Wu et al., 2014). Thus, we analyzed whether dysgenesis of the corpus callosum we detected in *crmp2^−/−^* mice is also accompanied by defects in callosal axon guidance. We *in utero* electroporated WT and *crmp2^−/−^* embryonal cortices with pCAGGS EGFP vector at E15.5, which results in labeling of cortical layer 2/3 (i.e. mainly callosal-projecting neurons). The brains were collected at postnatal day 6 (P6), fixed in 4% PFA and cut coronally to trace callosal axons in a hemisphere contralateral to the electroporation site (Fig. 2D, E). At this stage, axons from somatosensory cortex enter the contralateral cortex in WT (Wang et al., 2007). Most of the WT axons showed well organized, parallel growth. In contrast, *crmp2^−/−^* axons often failed to grow in an organized, parallel way upon leaving the main callosal tract, their distribution in the cortex seemed uneven. We tracked the electroporated axons after leaving callosal bundle at the contralateral site and quantified their tortuosity (i.e. the ratio of real length of the segment vs. distance of the first and last point of the segment, Fig. 2E). We found that tortuosity was significantly higher in *crmp2^−/−^* in comparison with WT (p<0.001, Fig. 2F).

Importantly, in some coronal sections of *crmp2^−/−^* mice we detected deregulated growth of callosal axons even in the midline. Together with the reduced length of corpus callosum in *crmp2^−/−^* mice in the rostro-caudal axis we detected (Fig. 2A – C) it was suggesting that CRMP2 deficiency may alter the rostro-caudal guidance of callosal axons in the midline. To test this hypothesis, we traced the callosal axons by injecting DiI into the fixed P9 and adult somatosensory cortices and analyzed organization and fasciculation of the traced axons in the midline (see Materials/Methods). We found that at P9, *crmp2^−/−^* axons were significantly more distorted as seen upon plotting to polar histograms or fan-in diagrams (interval (0°, ±20°): p=0.05, interval (±20°, ±40°): p=0.02, Fig. 2H – I, Fig. S3A, B).

Together, our results show that CRMP2 is an important regulator of axon growth and guidance in both peripheral (PNS) as well as central nervous system (CNS) and that out of the two major functions of CRMP2 (promotion of axon growth and mediation of Sema3A-dependent axon guidance) the mediation of axon guidance takes precedence over the growth promotion *in vivo*.

### CRMP2 mediates Sema3F-, but not Sema3A-driven axon pruning

CRMP2 has been associated with neurodevelopmental disorders like schizophrenia and ASD characterized by altered brain connectivity (Penzes et al., 2011) and defects in postnatal synaptic refinement through axon and dendrite pruning (Garey, 2010, Bourgeron, 2015). Importantly, in many regions Sema3F seems to play a more important role in pruning than Sema3A (Bagri et al., 2003, Low et al., 2008). Therefore, we asked whether CRMP2 regulates axon pruning and if, in this function, it mediates Sema3A or rather Sema3F signaling. For this, we analyzed two developing axonal systems showing either Sema3F-mediated pruning (the infrapyramidal bundle, IPB; (Bagri et al., 2003)) or Sema3A-mediated pruning (the hippocamposeptal bundle; (Bagri et al., 2003)). First, we analyzed stereotyped pruning of infrapyramidal bundle (IPB) of hippocampal mossy fibers taking place between P20 – P40 and regulated by Sema3F and its receptor complex (Nrp2/PlxnA3) (Bagri et al., 2003). At P14 (i.e. before pruning) calbindin immunostaining revealed presence of IPB in both WT as well as *crmp2^−/−^* mice (Fig. 3A, B). Synapses were formed in both IPB and the main bundles as revealed by VGluT1 staining (Fig. 3A, B). In 7 week old animals, however, when pruning was complete in WT, the IPB remained present in *crmp2^−/−^* mice, and their IPB index (IPB length/main bundle length) was significantly higher than in WT (p<0.001, Fig. 3C-G). The same pattern of IPB pruning was detected also in sagittal sections (Fig. S4A). To determine the maturity of IPB synapses, we stained adult coronal sections with antibodies against VgluT2 or VgluT1, which are both expressed in developing mossy fibers, but in adult hippocampus only VgluT1 is present (Liu et al., 2005). We found VgluT1 in the main bundles of WTs and *crmp2^−/−^* mice, as well as in the unpruned IPBs of *crmp2^−/−^* (Fig. 3E, F). We did not detect the immature VgluT2 signal (Fig. 3C, D). Thus, the IPB axons formed during postnatal development persist in *crmp2*^−/−^ mice into adulthood and form mature synapses. These results demonstrate that IPB pruning is defective in *crmp2^−/−^* mice (as is the case in *Sema3F^−/−^*, *Nrp2^−/−^* and *PlxnA3^−/−^* mice) (Bagri et al., 2003).

**Figure 3.**
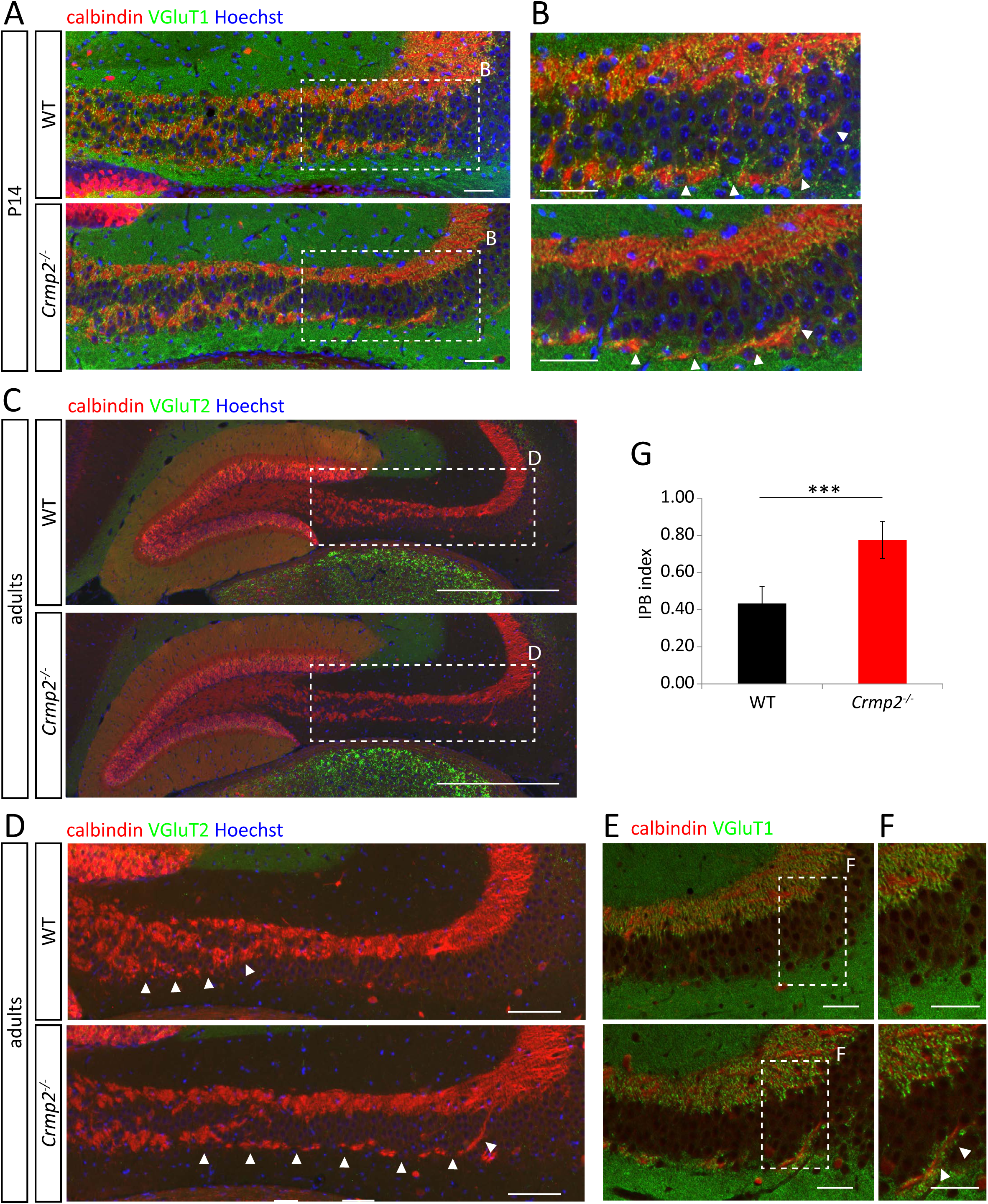
Infrapyramidal bundle fails to prune in *crmp2^−/−^* mice. (A) Coronal sections of P14 brains stained for calbindin and VgluT1. Infrapyramidal bundle (IPB) progresses into hippocampal CA3 region in both WT and *crmp2^−/−^* mice. (Counterstained with Hoechst) (B) Details from depicted regions in (A), arrows show the IPB. (C) Staining for calbindin shows unpruned infrapyramidal bundle in adult *crmp2^−/−^* mice. No VGluT2 signal in hippocampal CA3 region is present in WT and *crmp2*^−/−^. (D) Details from regions in (C) (arrows - IPB) (E, F) Staining for calbindin and VgluT1 shows mature synapses in the main bundle in both genotypes and in the IPB in *crmp2^−/−^* mice (arrows). (G) Quantification of IPB index (length of the IPB vs. main bundle, WT 0.43±0.05, *crmp2^−/−^* 0.78±0.05, p<0.001) counted from adult coronal sections (WT n=4, c*rmp2^−/−^* n=5), means± SD are shown. ***p<0.001, t-test. Scale bars: 100 μm in A and B, 500 μm in C, 100 μm in D, E and F. See also Fig. S4.

We next tested whether stereotyped pruning of a different group of axons arising from hippocampus – the hippocamposeptal axons – is also affected in *crmp2^−/−^* mice. CA1 neurons send their axons into medial and lateral septum at P0-1. However, at P8, only the axons to the lateral septum persist while the ones sent into the medial septum have disappeared (Linke et al., 1995). In this system, the pruning is mediated by Sema3A and not Sema3F (Bagri et al., 2003). Our analysis of the development of the hippocamposeptal axons in *crmp2^−/−^* mouse brains by means of tracers (Fig. S4B) shows no change in these brains as compared to WT brains. This indicates that the hippocamposeptal axon pruning is not affected by CRMP2 deficiency and that CRMP2 is a mediator of stereotyped axon pruning driven by Sema3F, but not Sema3A.

To further support the role of CRMP2 as a mediator of Sema3F-driven pruning, we analyzed pruning of corticospinal axons of visual cortex neurons that have previously been shown to be dependent on Sema3F signaling. In the early developmental stages, these neurons send their projections not only to the superior colliculus but also to two inappropriate targets, i.e. the inferior colliculus (IC) and the spinal cord. During the third postnatal week, inappropriate axons are eliminated through a pruning process regulated by Sema3F (Low et al., 2008). We analyzed the development of the visual cortex projection by the means of DiI anterograde tracing (Fig. 4A, B). In *crmp2^−/−^* mice, the inappropriate corticospinal axons were still largely present, and VP indexes (fluorescence intensity of corticospinal axons after vs. before the branch point) were not significantly different in *crmp2^−/−^* mice between adult and P9 stages (p=0.27, Fig. 4C), while they significantly dropped in WT (p<0.05, Fig. 4C). This data demonstrate that CRMP2 participates in postnatal refinement of corticospinal visual axons and is consistent with its role in mediating Sema3F signaling.

**Figure 4.**
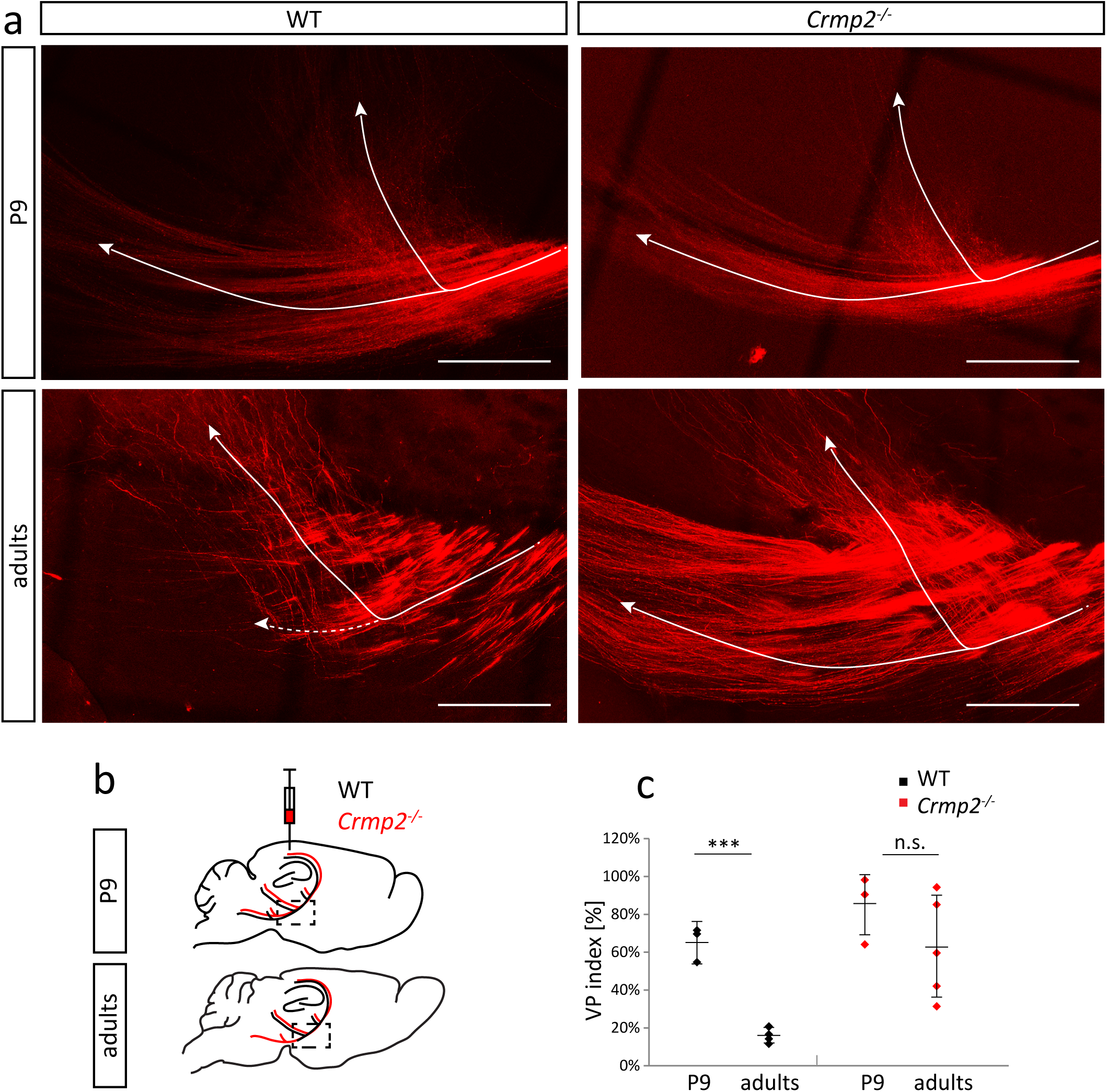
Defective pruning of corticospinal visual axons in *crmp2^−/−^* mice. (A) Upper row: DiI tracing of the visual cortex axons at P9 (before pruning, n=3), sagittal sections. Branching point of the tract is shown. Lower row: visual cortex axons in adult mice (n=5) after pruning period. Note significantly reduced number of axons continuing into pyramidal tract in WT. (arrows – 2 branches of corticospinal visual axons). Maximum projections are shown. (B) Schematic drawing of DiI injection and axon tracing. Axons that initially enter pyramidal tract fail to prune in *crmp2^−/−^* mice (red line). (C) Quantification of VP (visual pruning) index (fluorescence intensity of pyramidal axons after vs. before the branch point). Lower index thus indicates lower amount of axons in pyramidal tract. In WT animals after the pruning period, only a minor part of axons descends towards the pyramidal tract (P9 0.65±0.09, adults 0.17±0.04, p<0.001). However, in *crmp2^−/−^* mice, corticospinal axons are still largely present, and their VP indexes are not significantly different between adult and P9 stages (P9 0.54±0.18, adults 0.62±0.27, p=0.27). Means±SD are plotted. ***p<0.001, t-test. Scale bars: 200 μm. See also Fig. S5.

While in development, both CNS and PNS projections are refined by pruning, we did not detect changes in the pruning of the peripheral neuromuscular junctions (NMJs) in *crmp2^−/−^* mice using trigonum sterni muscle (the number of double-innervated NMJs was similar in WT and mutant mice at P11 (p>0.99, Fig. S4C, D)).

Finally, we asked if Sema3F-dependent axonal guidance is also mediated by CRMP2. Axon guidance is affected in two specific axonal bundles in Sema3F^−/−^ mice (Sahay et al. 2003) – namely the anterior commissure and the retroflex fascicle. We did not find any differences in formation of either of these axonal bundles in *crmp2^−/−^* mice (Fig. S5A, B) by immunostaining of P6 and P14 brains with anti-neurofilaments antibody (marker of axon tracts). Thus, CRMP2 seems to specifically mediate Sema3F-driven axon pruning but not Sema3F-driven axon guidance.

In conclusion, we found significant differences between WT and *crmp2^−/−^* mice in stereotyped pruning in regions controlled by Sema3F (visual cortex axons, IPB). In contrast we found no differences in regions controlled by Sema3A (i.e. hippocamposeptal axons) suggesting CRMP2 mediates Sema3F-driven, but not Sema3A-driven axon pruning.

### CRMP2 mediates Sema3F-dependent axon retraction in vitro

In order to directly show that CRMP2 is necessary for Sema3F-triggered axon pruning, but dispensable for Sema3A-dependent pruning, we tested pruning in an *in vitro* system. We prepared dissociated hippocampal cultures from WT and *crmp2^−/−^* E16.5 embryos and transfected them with EGFP at DIV1 (1 day in vitro). At DIV7, we added Sema3A or Sema3F into the cultures and analyzed the axon behavior in fluorescence microscope by time-lapse imaging. DIV7, which is an early stage of synapse formation (Basarsky et al., 1994), was chosen to facilitate analysis of neurons in still less complex connectivity patterns. (Fig. 5A, B, videos 2, 3). We analyzed only stable axon terminals that did not show any movement in one hour period prior to addition of the guidance cues (45% of all labeled axon terminals). In control conditions in both wild-type and knockout, a small number of the stable axons were spontaneously retracting (WT 13%, KO 20%, p=0.03, Fig. 5C). After addition of either Sema3A or Sema3F into WT culture, we observed a 3-fold increase in axon retractions (32% for Sema3A, p<0.001, 41% for Sema3F, p=0.0012, Fig. 5C). However, in *crmp2^−/−^*, this increase was detectable only after Sema3A (30%, p=0.0026) and not after Sema3F (22%, p=0.23). This data demonstrate, that in primary neuron cultures undergoing synaptogenesis, CRMP2 is essential to mediate Sema3F but not Sema3A signaling. This is in agreement with our *in vivo* findings that stereotyped pruning in *crmp2^−/−^* is affected in Sema3F-controled, but not in Sema3A-controled regions.

**Figure 5.**
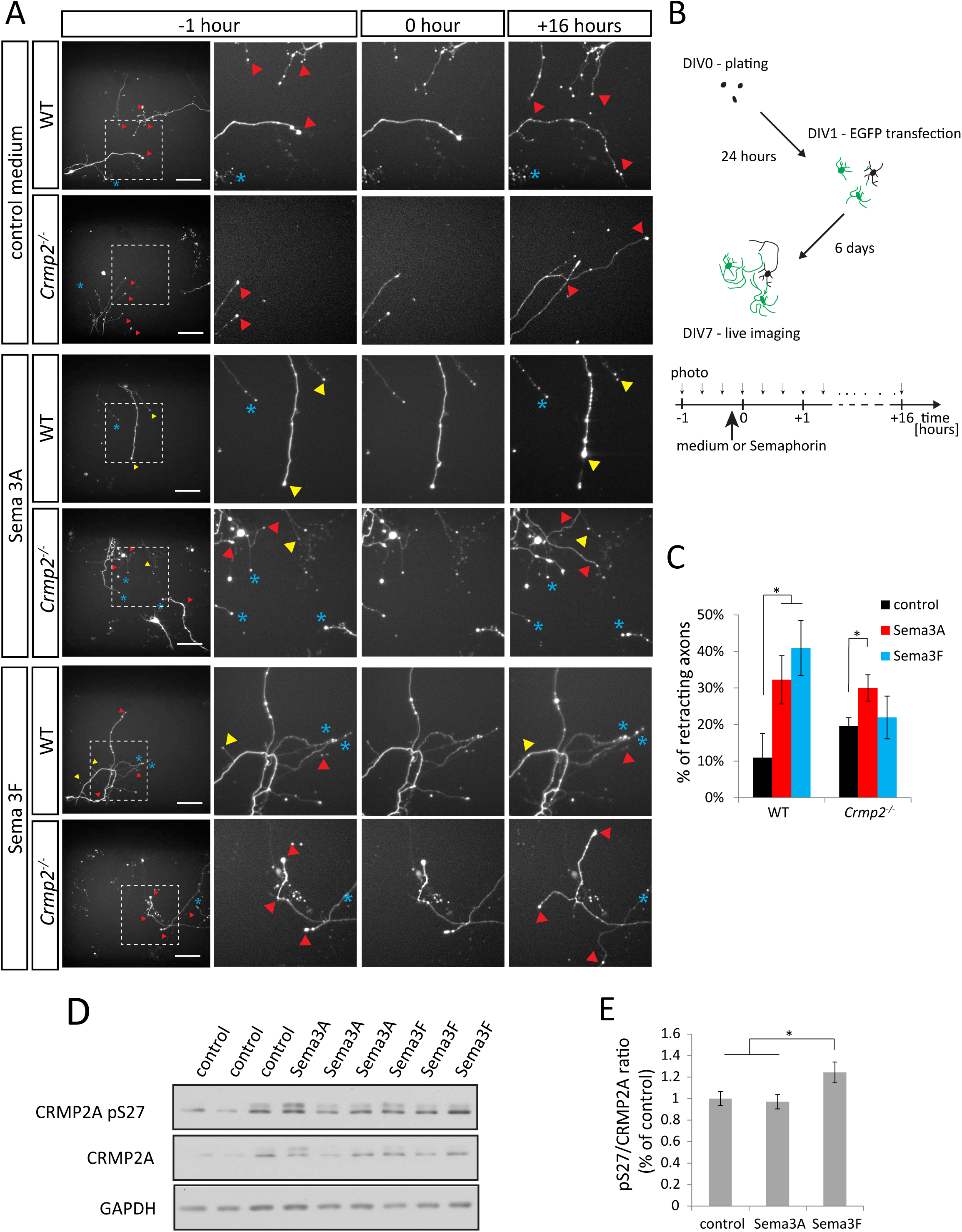
CRMP2 mediates Sema3F signaling in primary neurons. (A) Live imaging (16h) of DIV7 cultured hippocampal neurons to assess axon retraction/pruning after Semaphorin stimulation. Upper panel: Axon behavior in medium without Semaphorins. Middle panel: stimulation with Sema3A (1nM, n=669 axons for WT, 761 axons for knockout) causes retraction of both WT and *crmp2^−/−^* neurons. Lower panel: stimulation with Sema3F (5 nM, n=602 axons for WT, 955 axons for knockout) causes axon retraction in WT, but not in *crmp2^−/−^* neurons. Red triangles show growing axons, yellow show retracting axons and blue asterisks indicate steady non-growing axons. See also supplementary material videos (Video 2, 3). (B) Schematic drawing of the experimental setup. (C) Quantification of retracting axons (number of retracting vs. steady axons, 3 experiments; WT, control: 13.4±5%, Sema3A: 31.2±6%, p<0.001, Sema3F 36.9±8.7%, p<0.05; *crmp2^−/−^*, 19.8±1.8%, 28.5±4.4% p<0.05, 22.4±4.1%, p=0.23), means±SD are shown. (D) Western blot analysis of CRMP2A phosphorylation at Ser27 in DIV4 hippocampal neuronal lysates after class 3 Semaphorin stimulation. (E) Quantification of the western blots. Sema3F increases ratio of CRMP2A pS27 (normalized to total CRMP2A). Means±SEM are present in E. *p<0.05. Scale bars: 100 μm.

Finally, we asked if Sema3F acts on CRMP2 by inducing changes in its phosphorylation status. We isolated hippocampal neurons from E18.5 rat embryos (to increase neuron yield), treated them at DIV4 with Sema3A or Sema3F for 3 hours and analyzed the levels of Ser27 phosphorylation in the cell lysates by western blotting (Ser27 is a CDK5 phosphorylation site (Balastik et al., 2015) at CRMP2A isoform, which we found highly phosphorylated in hippocampus (not shown)). Ratio of pS27-CRMP2A/CRMP2A was increased only after Sema3F stimulation (p<0.05, Fig. 5D, E), indicating that phosphorylation of Ser27 is downstream of Sema3F stimulation.

### CRMP2 regulates dendritic spine remodeling in hippocampal granule cells

Besides triggering axon pruning, Sema3F regulates also the development of some classes of dendritic spines (e.g. spines of dentate gyrus (DG) granule cells) (Tran et al., 2009). In contrast, Sema3A/Nrp1 signaling seems to be dispensable for dendritic spine morphogenesis (Tran et al., 2009, Yamashita et al., 2007). In order to test whether CRMP2 participates also in spine development/morphogenesis, we diolistically labeled DG neurons and analyzed their dendritic spines. Surprisingly, we found significantly increased spine density in *crmp2^−/−^* adult DG granule cells compared to WTs (Fig. 6B, C, p<0.001). This phenotype was similar to that found in *Sema3F^−/−^* and *Nrp2^−/−^* (Tran et al., 2009). Next, we analyzed branching of DG granule cell dendrites. Sholl analysis revealed no differences between WT and *crmp2^−/^*^−^ mice (Fig. 6D, E, p>0.99), which is again in line with the phenotype of *Sema3F^−/−^* mice (Tran et al., 2009).

**Figure 6.**
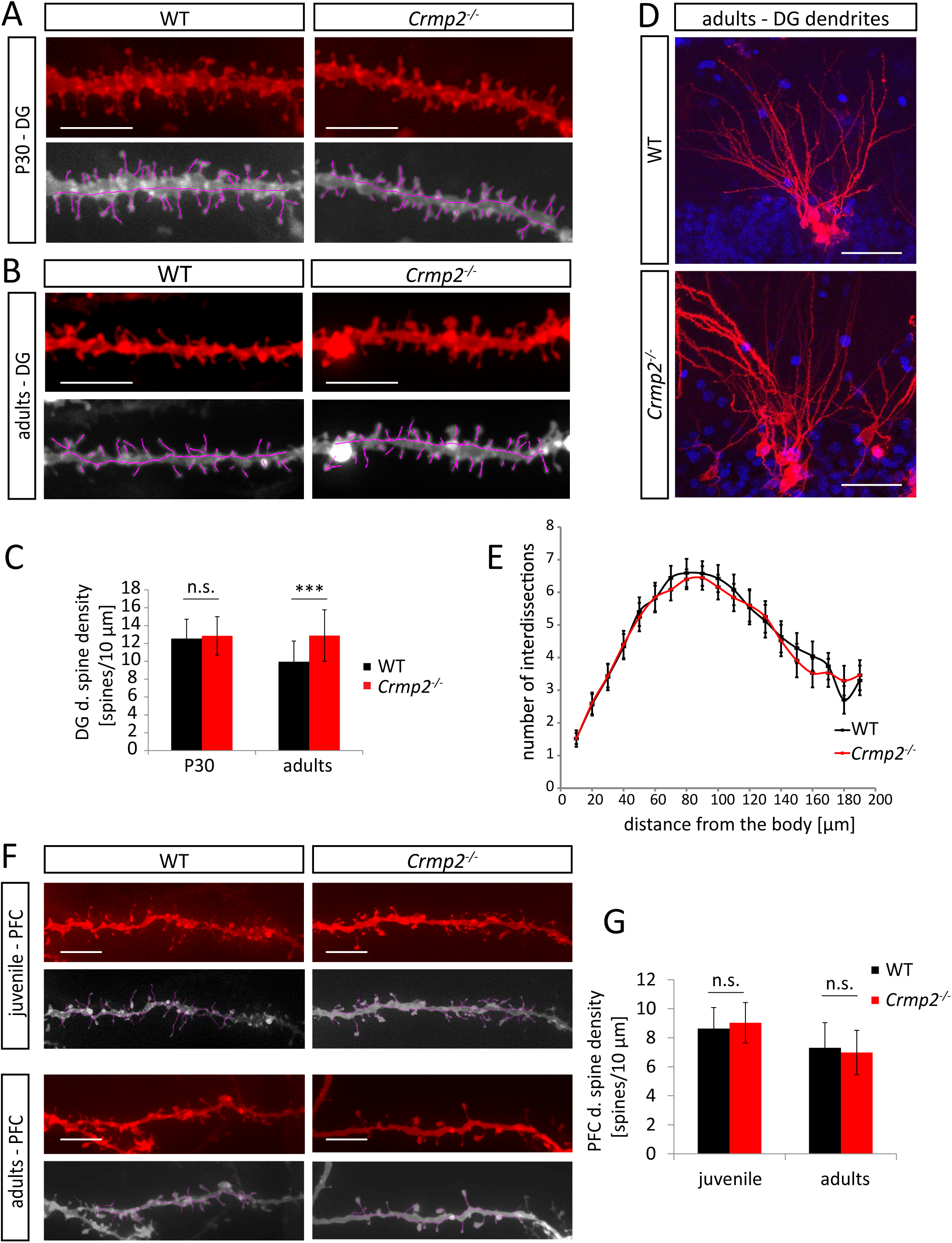
CRMP2 regulates dendritic spine refinement in dentate gyrus granule cells. (A) Dendritic spine density in DiOlistically labeled DG granule cells (n=3 animals, WT=61 dendrites, knockout=64 dendrites) is similar in WT and *crmp2^−/−^* mice at P30. (B) Spine density is reduced in adult WT but no *crmp2^−/−^* granule cells (n=3, WT=41 dendrites, knockout=37 dendrites). (C) Quantification of dendritic spine density in DG granule cells 50 – 100 μm away from the soma (WT, P30: 12.5±2.2 spines/10 μm, adults: 9.95±2.3; *crmp2^−/−^*, P30: 12.86±2.1 p=0.47, adults: 12.88±2.9 p<0.001) means±SD are shown (D) Analysis of branching of DiOlistically labeled DG granule cell dendrites in adult WT and *crmp2^−/^*^−^ mice (n=3 animals, ≥25 dendrites). (E). Quantification of granule cell branching by Sholl analysis (p>0.99), means±SEM are shown. (F) Spine density in prefrontal cortex (PFC) is similar in WT and *crmp2^−/^*^−^ mice in both juvenile (P25, n=3, ≥50 dendrites) and adult mice (n=3, ≥50 dendrites). (G) Quantification of dendritic spine density of PFC pyramidal neurons, basal dendrites (WT, juvenile: 8.63±1.44 spines/10 μm, adults: 7.3±1.4; *crmp2^−/−^*, P30: 9.05±1.73, p=0.18, adults: 6.98±1.53, p=0.25), means±SD are shown. ***p<0.001, t-test (C, G), ANOVA (E). Scale bars: 5 μm in A, B, F, 50 μm in D. See also Fig. S6.

Higher spine density could be a result of either increased generation of new spines, or defective pruning of spines, or both. Considering the axon pruning defects we found in Sema3F-regulated areas in *crmp2^−/−^* mice (see above), and considering Sema3F promotes loss of spines *in vitro* (Tran et al., 2009), we hypothesized that Sema3F regulates dendritic spine pruning through CRMP2. To test this hypothesis, we labeled and counted DG dendritic spine density in P30 (adolescent) mice when dendritic spines are virtually all formed and the pruning process starts (Petanjek et al., 2011, Bian et al., 2015). We found no differences in dendritic spine density between WTs and mutants at P30 (Fig. 6A, C, p=0.47). This indicates that the formation of spines is unaltered in *crmp2^−/−^* mice and that it is the process of dendritic spine pruning that is defective in these mice. As DG spine morphogenesis is regulated through Sema3F (Tran et al., 2009), our findings indicate that CRMP2 mediates Sema3F-dependent pruning also in dendritic spines.

Defects in the distribution of dendritic spines due to aberrant synapse refinement are one of the key features of both ASD and schizophrenia. Generally, dendritic spine number in ASD patients is higher than in control subjects or variable in different regions, while in schizophrenia patients it is lower (Penzes et al., 2011). This applies particularly to some brain regions, e.g. prefrontal cortex (PFC), where excessive spine elimination has been associated with pathogenesis of schizophrenia (Garey et al., 1998). Since some phenotypical aspects of conditional *crmp2^−/−^* mice like impaired sensorimotor gating have been related to schizophrenia (Zhang et al., 2016), we asked whether CRMP2 deficiency also leads to spine overpruning and reduced spine density in PFC. At P25, PFC spine density was similar in both WT and mutants (Fig. 6F, G, p=0.18) and similar to our findings in the DG. Importantly, unlike in DG, we did not detect any significant difference in PFC spine density between adult WT and *crmp2^−/−^* mice (Fig. 6F, G, p=0.25), which is not consistent with schizophrenia-like phenotype.

### Crmp2^−/−^ mice display juvenile sociability defects, memory impairment and decreased anxiety

While schizophrenia shares some behavioral symptoms with autism (e.g. cognitive and social deficits), it differs in the onset of the disease, as ASD manifests typically in 3-years old children while schizophrenia in the adolescence at the earliest (Lord et al., 2018, Garey, 2010). This suggests that any changes underlying ASD must be present already very early in postnatal development. To assess this hypothesis, we wanted to see whether CRMP2 deficiency results in juvenile behavioral changes. To this effect, we analyzed ultrasonic vocalization in P6, P8 and P12 WT and *crmp2^−/−^* pups as a measure of their sociability (Ey et al., 2013). We recorded vocalization of WT and mutant pups in 5-minutes intervals after isolation from their mothers and found that at P6 the number of calls was similar between both groups (p=0.73), but decreased significantly in the mutants at P8 (p=0.015) (Fig. 7A – C). At P12, the mutants were almost completely silent (p=0.03); in fact, only two mutant pups from 13 vocalized at all (10/14 in WT). The duration of the individual calls was also significantly shorter in *crmp2^−/−^* pups in both P8 (p=0.011) and P12 (p=0.03) (Fig. 7B). This early onset of defects in social behavior has been described in numerous mouse models of ASD (Bourgeron, 2015).

**Figure 7.**
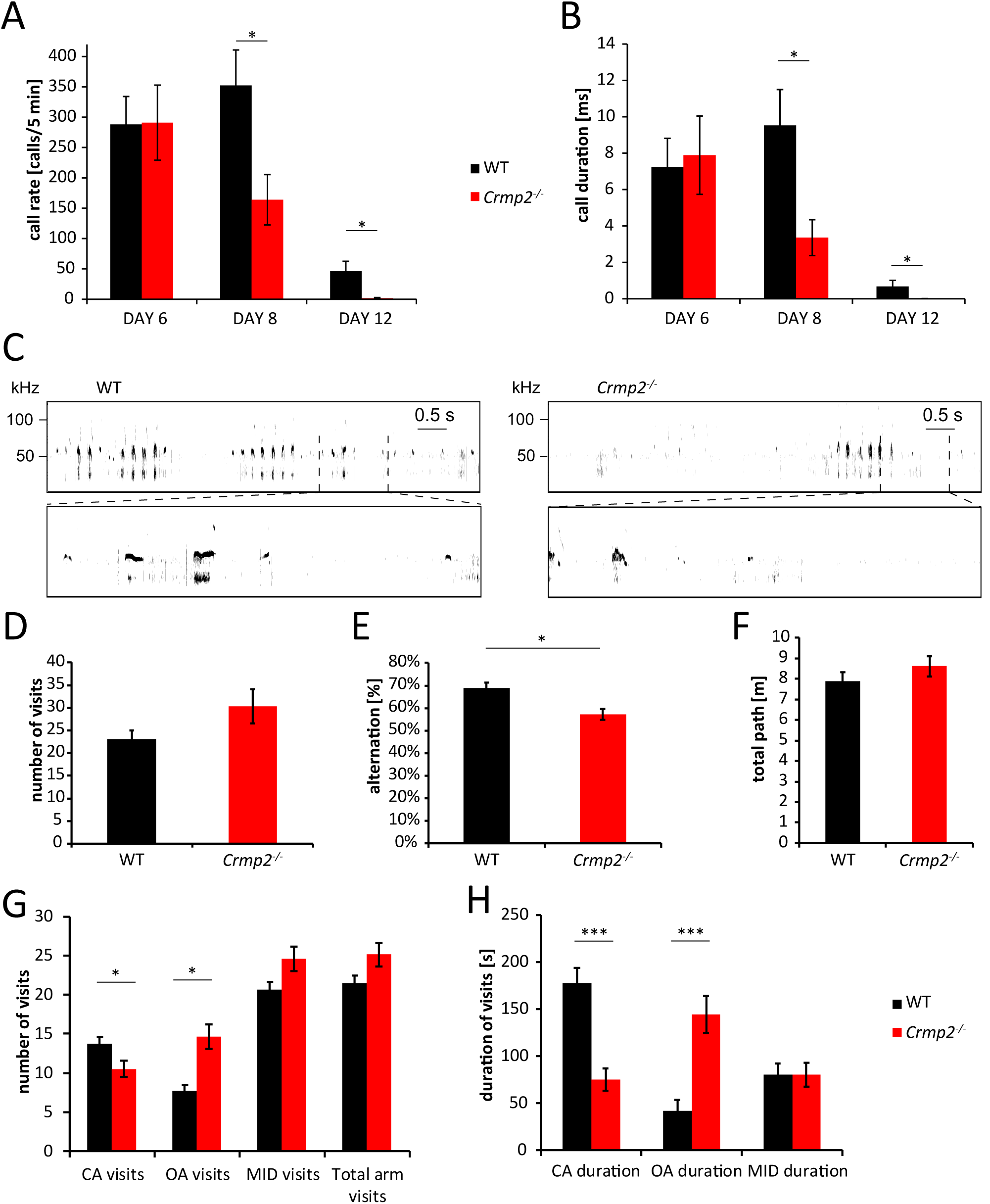
*crmp2^−/^*^−^ mice show early onset behavioral defects. (A – C) Ultrasonic vocalization was measured at P6, P8 and P12 (WT n=14, *crmp2^−/−^* n=13). In *crmp2^−/−^* mice, there is a significant decrease in the rate and duration of calls at P8 (call rate, WT 352±58/5 min, *crmp2^−/−^* 164±41/5 min, p=0.015; call duration, WT 9.52±2 ms, *crmp2^−/−^* 3.36±1 ms, p=0.011) and P12 (call rate, WT 46±16/5 min *crmp2^−/−^* 1.54±1.24/5 min, p=0.03; call duration, WT 0.68±0.35 ms, *crmp2^−/−^* 0.007±0.006 ms, p=0.03), means±SEM are shown. (C) Representative sonograms of the P8 mice. (D, E) Y maze test (WT n=9, *crmp2^−/−^* n=8). Decreased alternations between arms of the maze suggest impaired working memory (visits, WT: 23.1±1.9, *crmp2^−/−^* 30.4±3.8, p=0.12; alternations, WT 38.8±2.4%, *crmp2^−/−^* 57.3±2.4%, p=0.006), means±SEM are shown. (F – H) Elevated plus maze test (n=10). Total distance walked is similar in WT and *crmp2^−/−^* (WT 7.9±0.45 m, *crmp2^−/−^* 8.6±0.5 m, p=0.3). Frequency and duration of open arm (OA) visits is increased in *crmp2^−/^*^−^ mice suggesting decreased anxiety (CA frequency: WT 13.7±0.9/5 min, *crmp2^−/−^* 10.5±1.1/5 min, p=0.04; OA frequency: WT 7.7±0.8/5 min, *crmp2^−/−^* 14.6±1.7/5 min, p=0.002; CA duration: WT 177±17 s., *crmp2^−/−^* 75.3±12.6 s, p<0.001; OA duration: WT 42±12.6 s, *crmp2^−/−^* 144.4±21 s, p<0.001), means±SEM are shown. *p<0.05,***p<0.001, t-test or Kruskal-Wallis (P12 vocalization). See also Fig. S6.

We next asked whether hippocampus-dependent memory functions are affected in *crmp2^−/−^* mice as they exhibit aberrant inputs from DG into CA3 (unpruned mossy fibers) and increased spine density in DG granule cells (input from entorhinal cortex). We tested working memory using Y-maze (a three-arm maze) where WT and mutants showed comparable level of exploratory activity (Fig. 7D). However, the ratio of spontaneous arm alternation was significantly lower in *crmp2^−/−^* mice (Fig. 7E, p=0.0058) indicating a working memory impairment (Zhang et al., 2016). In contrast, long-term memory and general behavioral flexibility seem not to be affected in *crmp2^−/−^* mice as revealed by active place avoidance on a rotating arena test (Fig. S6C).

Similar as the other CRMP2 deficient mouse models (Zhang et al., 2016, Nakamura et al., 2016), we detected anxiety impairment in *crmp2^−/−^* mice using elevated-plus maze. In the task, the knockouts also demonstrated increased activity during exploration of the maze (Fig. 7F), spent more time in the open arms of the maze (p<0.001) and visited them more often (p=0.0016) than their WT counterparts (Fig. 7 G, H) suggesting decreased anxiety, or perhaps a more general lack of adequate response to potentially dangerous situations. This phenotype may reflect the hippocampal phenotype of *crmp2^−/−^* mice as lesions in particularly ventral hippocampus have been shown to result in similar decreased anxiety in mouse models (Kjelstrup et al., 2002).

## DISCUSSION

CRMP2 has been long considered an important regulator of Semaphorin 3A-mediated axon guidance during embryonic development. Its expression, though, is high even in the early postnatal neurons, but its role in the postnatal development and adult neurons has so far been elusive. In the present study we demonstrated that CRMP2 is not only mediator of Sema3A signaling regulating axon guidance in embryonic development, but importantly, that it plays a central role in the postnatal refinement of the nervous system. By generating new *crmp2^−/−^* mice and analyzing their phenotype we first showed that CRMP2 deficiency *in vivo* leads to axon guidance defects in CNS and PNS that could be attributed to changes of Sema3A signaling. Strikingly, we demonstrated that CRMP2 mediates also Sema3F signaling and that CRMP2 deficiency disrupts early postnatal Sema3F-mediated axon and dendritic spine refinement in multiple areas of the CNS. Changes in Sema3F signaling pathway have been considered a risk factor in the pathogenesis of ASD. In accord with that, we showed that *crmp2^−/−^* mice suffer from altered pruning and early postnatal social interaction defects previously linked to autism. Together, our *in vivo* and *in vitro* data demonstrated a novel function of CRMP2 in postnatal fine-tuning of the nervous system by Sema3F, and showed that its deficiency in mice leads to neurodevelopmental defects associated with pathogenesis of the ASD in human.

### Regulation of axon and dendritic pruning by Sema3F and CRMP2

Initial growth of axons and dendrites during embryonic and early postnatal period results in an embryonic template that must be later refined to generate a functional healthy nervous system. (Luo and O’Leary, 2005). Stereotyped refinement of nervous system was uncovered in several regions: (a) Infrapyramidal bundle (IPB) axons of hippocampal mossy fibers are retracted between P14 – P30 in mice (Riccomagno et al., 2012); (b) Excess of layer 2/3 callosal axons of motor, sensory and visual cortex is refined until P30 (Zhou et al., 2013); (c) Corticospinal axons from layer 5 visual cortex are eliminated between P9 – P25 (Low et al., 2008). In addition, density of dendritic spines in various brain regions in mice and humans peaks between childhood and adolescence (Bian et al., 2015, Petanjek et al., 2011) and its significant portion is eliminated until adulthood. Finally, pruning also occurs in peripheral nervous system – originally polyneuronal innervation of muscle fibers is later in development refined so that each muscle fiber is innervated by only one motor axon (Tapia et al., 2012, Brill et al., 2016).

Knockout mice studies identified extracellular cues and receptors that mediates pruning in rodents. These include Sema3A and its receptor complex (Nrp1/PlxnA4), and Sema3F and its receptors (Nrp2/PlxnA3), EphrinB3-EphB2 reverse signaling and C4 component of complement (Bagri et al., 2003, Xu and Henkemeyer, 2009, Cocchi et al., 2016). Intracellular pathways that translate extracellular signal to cytoskeleton are however poorly understood. Here, we identified CRMP2 as a novel mediator of pruning in rodent brain triggered by Sema3F.

Sema3F binds preferentially to Nrp2/PlxnA3 receptor complex and is important regulator of neural development. Deficiency of Sema3F, Nrp2 or PlxnA3 results in defects in stereotyped pruning of hippocampal infrapyramidal bundle, distribution of DG dendritic spines, or anterior commissures, (Riccomagno et al., 2012, Tran et al., 2009). Moreover, *PlxnA3/A4^−/−^* mice display defects in pruning of visual axons. Similarly, in our *crmp2^−/−^* mice, we found all: IPB pruning defect, alteration of DG spine density (Figs 3, 6) as well as defects in pruning of visual cortex axons (Fig. 4). Moreover, our *in vitro* assays (Fig. 5) showed Sema3F is unable to induce axon retraction in hippocampal neurons isolated from *crmp2^−/−^* embryos.

Next to Sema3F, Sema3A has also been shown to orchestrate pruning of e.g. hippocamposeptal or callosal axons (Vanderhaeghen and Cheng, 2010, Zhou et al., 2013). As CRMP2 has originally been identified as a mediator of Sema3A (Goshima et al., 1995), we hypothesized that CRMP2 is involved also in Sema3A-triggered pruning. However, we didn’t find any evidence that would support this hypothesis. We did not detect hippocamposeptal axons by retrograde tracing after the pruning period in *crmp2^−/−^* pups (Bagri et al., 2003) (Fig. S4B). In callosal axons, we found significant differences in axon guidance between WT and *crmp2^−/−^* mice in P6 – P9 mice (Fig. 2), but the significance was lost in adult mice (Supp. Fig. 3C – E), which could be due to presence of effective Sema3A-dependent axon pruning. In line with these *in vivo* data, we found that Sema3A (but not Sema3F) was able to partially induce axon retraction of the stalling *crmp2^−/−^* axons in 1-week-old hippocampal neuron cultures (Fig. 6). It is possible that other members of CRMP family partially rescue CRMP2 deficiency in Sema3A signaling. This would be in agreement with the only mild defects we detected in *crmp2^−/−^* peripheral nerves (Fig. 1C, D).

Importantly, we also did not find any defect in the pruning of neuromuscular junctions (NMJs) in *crmp2^−/−^* mice (Fig. S4C, D). At the end of the embryonic development each synapse is innervated by up to ten axon branches of different motor units (Tapia et al., 2012). During the first two postnatal weeks, all except one terminal branch are pruned back establishing singly innervated NMJs (Sanes and Lichtman, 1999). While the exact molecular cascade regulating motor axon pruning is not known, Sema3A seems to play a role in the process as its receptor, Nrp1, is expressed in presynaptic axon terminals (Venkova et al., 2014). Moreover, Sema3A secreted from Schwann cells participates in NMJ remodeling (De Winter et al., 2006). Sema3F signaling has so far not been linked to motor axon pruning.

Dendritic spine density changes dynamically during childhood and adolescence. In mice, spine density peaks around 1 month and then decreases to reach stable levels around 2 months (Bian et al., 2015). It has been shown that distribution of dendritic spines is regulated by class 3 Semaphorins (Tran et al., 2009). Previous *in vitro* experiments showed that Sema3F, but not Sema3A, decreases PSD-95 positive puncta in dissociated DG neurons (Tran et al., 2009). Accordingly, Sema3F-treated cortical neurons displayed decrease in apical dendrite spine density (Tran et al., 2009). Adult *Sema3F^−/−^*, *Nrp2^−/−^* and *PlxnA3^−/−^* mice show increased spine density in several brain regions, in particular DG dendrites (Tran et al., 2009, Demyanenko et al., 2014). *Crmp2^−/−^* mice mimic this phenotype as we also found increased spine density in adult DG granule cells (Fig. 6). Interestingly, in *Sema3F^−/−^* mice, increased DG spine density is detectable already during spine generation (P21) and is largely retained into adulthood, while in WT they are subsequently pruned (Tran et al., 2009). In *crmp2^−/−^ mice*, we did not find increased DG spine density in the pre-pruning period (P30, Fig. 6A), but similar to Sema3F^−/−^ mice, we found defects in DG spine pruning. Together, this data suggest that while Sema3F signaling regulates both spine generation and pruning, CRMP2 controls only the spine pruning.

### CRMP2 in axonal growth in vivo

As demonstrated in knockout lines of Sema3A and its downstream targets (*Sema3A^−/−^*, *Nrp1^−/−^*, *PlxnA4^−/−^* mice), Sema3A signaling is an essential regulator of development of rodent trigeminal nerve, facial nerve, DRGs projection, olphactory bulb, hippocampal formation and corpus callosum (Kitsukawa et al., 1997, Taniguchi et al., 1997, Schwarting et al., 2000, Friedel et al., 2005, Pozas et al., 2001, Schwarz et al., 2008, Zhou et al., 2013, Wu et al., 2014). Surprisingly, although Sema3A or Nrp1 deletion causes strong overgrowth of some peripheral nerves (e.g. trigeminal and spinal axons) (Kitsukawa et al., 1997, Taniguchi et al., 1997), we found only a mild overgrowth and increased branching of these axons upon deletion of its downstream mediator CRMP2 in *crmp2^−/−^* mice (Fig. 1, Fig. S2). This could be due to a partial rescue of Sema3A signaling in these neurons by other CRMP family members as mentioned before.

Electroporation studies of *Sema3A^−/−^* or *Nrp1^floxed/floxed^* brains demonstrated their role in the development of corpus callosum, with mispositioned axons in callosal midline and axonal mistargeting in contralateral cortex at P8 (Zhou et al., 2013). Defects in axon pruning of this region were also suggested (Zhou et al., 2013). Using the in utero electroporation and DiI tracing in *crmp2^−/−^* mice we also found defective guidance of callosal axons in the contralateral cortex and their altered positioning and orientation in the midline in the rostrocaudal axis (Fig. 2).

Notably, Sema3F signaling is essential also for guidance of specific cranial nerves and was related to development of limbic system and anterior commissure (Sahay et al., 2003, Giger et al., 2000). CRMP2 seems to be dispensable for Sema3F-mediated axon guidance in these regions as we found no major disruption of axon tracts guided by Sema3F in *crmp2^−/−^* mice (Fig. S5).

Previous *in vitro* and *in vivo* experiments suggested that CRMP2 also regulates neuronal migration (Ip et al., 2014). However, using the *in utero* electroporation, we found no significant changes in neuron distribution in the developing WT and mutant cortical plates at E17.5 (Fig. S1F). This likely reflects different experimental paradigms used in the studies (somatic knockdown vs. full knockout) (Ip et al., 2014).

### CRMP2 involvement in pathogenesis of neurodevelopmental disorders

Deregulation of CRMP2 has been linked to several neurodevelopmental disorders (Sfari Gene database, https://gene.sfari.org/database/human-gene/DPYSL2). Recently published analysis of conditional brain-specific (Zhang et al., 2016) and full *crmp2* knockout mice (Nakamura et al., 2016) showed multiple behavioral defects associated with CRMP2 deficiency. Notably, conditional knockout mice revealed hyperactivity and prepulsed inhibition (PPI) deficit together with social behavior impairment. PPI serves as a test for evaluating sensorimotor gating – the phenomenon that is often altered in schizophrenia patients. In addition, clozapine (an antipsychotic drug) treatment was capable to reduce hyperactivity in conditional *crmp2^−/−^* mice. Furthermore, morphological analysis showed increased volume of brain ventricles and impaired dendritic development in hippocampal CA1 and DG neurons, which is associated with schizophrenia and other neurodevelopmental disorders (Zhang et al., 2016). Analysis of the two CRMP2 mouse knockout models revealed also their significant differences. In particular, while PPI was reduced in the conditional mice, in the full *crmp2^−/−^* mice it was not significantly different to WT (Zhang et al., 2016, Nakamura et al., 2016). This suggests that even a minor difference in the spatio-temporal inactivation of CRMP2 during development can have a major impact on the development and severity of the resulting neurodevelopmental defects. The analysis of the full CRMP2 knockout mice, which we generated, revealed several phenotypical defects present in the conditional and full CRMP2 knockout mice (e.g. ventriculomegaly, spine density changes in DG, working memory defects, or hyperactivity) (Zhang et al., 2016, Tobe et al., 2017). In addition, though, we found that CRMP2 knockout leads to axonal pruning and dendritic spine remodeling defects compatible with ASD rather than schizophrenia (Lord et al., 2018) (Fig. 3, 4, 6). These changes were accompanied by altered social communication in early postnatal (P8 and P12) mutants (Fig. 7) further corroborating the role of CRMP2 in the pathogenesis of ASD.

Importantly, Sema3F signaling has been also implicated in the pathogenesis of ASD. Sema3F or NRP2 deficient mice show both behavioral and neuropathological aspects of ASD (Li et al., 2019) and Sema3F interacts with multiple ASD-related genes e.g. fragile X mental retardation protein or MECP2 (Degano et al., 2009). Thus, by linking Sema3F and CRMP2 signaling and comparing the histological as well as behavioral effects of their deficiency, our data strongly implicate that the Sema3F-CRMP2 signaling plays an important role in ASD pathogenesis. Since previously CRMP2 has been linked with pathogenesis of schizophrenia, it may serve as a molecular link linking class 3 Semaphorin signaling defects in both autism and schizophrenia.

## METHODS

### Animals

All animal studies were ethically reviewed and performed in accordance with European directive 2010/63/EU and were approved by the Czech Central Commission for Animal Welfare. Mice, all in C57BL6/N background, were housed and handled according to the institutional committee guidelines with free access to food and water. Unless stated otherwise, adult mice used for experiments were 12 – 16 weeks old. See table S1 for number of animals used in experiments.

### Crmp2^−/−^ mice generation

We used TALEN mutagenesis (transcription activator-like effector nucleases) to generate *crmp2^−/−^* mice. Two TALEN pairs targeting sequences 185-150bp 5’ of *crmp2* exon 2 and 183-218bp 3’ of exon 3 (Fig. 1a) were designed using TAL Effector Nucleotide Targeter 2.0 (https://tale-nt.cac.cornell.edu/) (Cermak et al., 2011, Doyle et al., 2012), assembled using the Golden Gate Cloning system (Cermak et al., 2011), and cloned into the ELD-KKR backbone plasmid. DNA binding domains of TALENs specific for the desired target sites within the crmp2 locus consisted of following repeats: HD-HD-NN-HD-HD-HD-NG-NI-NN-HD-NG-NN-NN-NI-NG-HD-NG (5’ TALEN-crmp2-ex1), NN-HD-NI NI-NG-HD-HD-NG-HD-NG-NN-NG-HD-NG-HD-NG-NG (3’ TALEN-crmp2-ex1) HD-HD-NI-NI-NN-NI-NN-NG-HD-NI-HD-NG-NN-NI-NN-HD-NG-NG (5’ TALEN-crmp2-ex2) and NN-HD-NI-HD-NI-NG-NG-HD-NG-NI-HD-HD-NI-NI-NG-NN-NG (3’ TALEN-crmp2-ex2). All TALEN plasmids were used for production of TALEN encoding mRNA as described previously (Kasparek et al., 2014). TALEN mRNAs (with total RNA concentration of 40 ng/μl) were microinjected into C57BL6/N-derived zygotes. Genomic DNA isolated from tail biopsies of newborn mice was screened by PCR for deletion of exon1 and 2 (3673 bp) (primers F1: 5’-ATATCCCACGATTCTGACCAATCA-3’ and R1: 5’-CCAAATAACTGCAGTGTAGCCTAT-3’), deletion was confirmed by locus sequencing and mice used as founders of *crmp2^−/−^* line. The mouse line was genotyped by PCR using locus specific primers: R1: ACTTACCGTGATGCGTGGAA, F1: TCACCCTCCCGGGACGAT, R2: TCTACCAATGTTACAACACAGA.

### Antibodies, cell dyes and plasmids

Primary antibodies used in this study are as follows: mouse anti CRMP2 (WAKO, IHC 1:200, WB 1:5000), rabbit affinity purified anti CRMP2A (IHC 1:75) (Balastik et al., 2015), rabbit anti CRMP2A (WB, 1:30 000) (Balastik et al., 2015), mouse anti neurofilaments (2H3 antigen, DSHB Iowa, 1:150), rabbit anti MAP2 (Abcam, 1:300), rabbit anti calbindin (Swant, 1:600), mouse anti VgluT2 and VGluT1 (Millipore, 1:400), rabbit anti pCRMP2A-S27 (phospho-specific polyclonal antibodies generated in rabbit using S27-phospho-peptide CNLGSG(Sp)PKPRQK and affinity purified in two steps against S27-non-phospho and S27-phospho-peptides, IHC 1:1200, WB 1:20000), mouse anti tau (Abcam, 1:500), mouse anti-βIII-Tubulin conjugated to Alexa 488 (BioLegend, 1:200). Secondary fluorescent antibodies were conjugated with various Alexafluor dyes: anti mouse (Alexa 488), anti rabbit (Alexa 594), diluted 1:400. Hoechst 33342 (1 μg/ml) was used to counterstain cell nuclei. In some cases, ABC kit (Vector laboratories) was used for detection. For whole mount immunostaining, secondary anti mouse antibody conjugated with HRP was used (1:1000). For western blots, secondary anti mouse or anti rabbit conjugated with HRP were used (1:10 000). Choleratoxin subunit B conjugated to Alexa 647, DiI and DiO were purchased from Thermo Fisher. α-bungarotoxin conjugated to Alexa 594 was purchased from Invitrogen (50 µg/ml; 1:50). EGFP was cloned into pCAGGS vector.

### Histology, immunohistochemistry, biochemistry

Mice were perfused transcardially with PBS and ice-cold 4% paraformaldehyde (PFA) in PBS. Brains were isolated and postfixed overnight at 4°C in 4% PFA/PBS. Subsequently, brains were washed in PBS and processed as described previously (Balastik et al., 2015). 7-μm-thick paraffin sections were created. Immunohistochemistry was done as described previously (Balastik et al., 2015). Sections were deparaffinated as follows: 2x 10 min. 100 % xylene, 2x 10 min. 100% ethanol, 3 min. 90% EtOH, 3min. 70% EtOH, 3 min. 50% EtOH, PBS. Antigen retrieval was performed in some cases using citrate-based antigen retrieval solution (Vector) diluted 1:100 in water. Slices were blocked in 1% BSA/0,2% Tween/PBS (PBST) and incubated with primary antibodies overnight at 4°C. Next, slices were washed 3x 5 min. in PBST and incubated with secondary antibodies conjugated with alexa fluors, 2 hours at RT. Then, slices were washed 3x 5 min. in PBST and mounted in Mowiol with Hoechst (1:1000). For bright field microscopy, slices were pretreated 15 min. in 3% H_2_O_2_ prior to blocking and ABC kit (Vector) was used according manufactures instructions. HRP activity was detected with 0.05% DAB. Subsequently, slices were dehydrated in ethanol – xylene and embedded into Eukitt. For biochemical analysis of CRMP2 phosphorylation, rat E18.5 neurons were plated in 3.5 cm Petri dishes coated with laminin (1 μg/ml) and poly-D-lysine (50 μg/ml). Lysates were prepared at DIV4 after 3-hours incubation with Sema3A (1 nM) or Sema3F (5 nM). SDS-PAGE and western blotting were done as described previously (Balastik et al., 2015).

### Whole-mount immunohistochemistry

Whole-mount immunohistochemistry was performed as described previously (Balastik et al., 2015) with some modifications. E10,5 – E12,5 embryos were isolated from time-pregnant mothers. WTs and knockouts from the same litter were compared. Embryos were fixed in 4% PFA/PBS overnight, 4°C. Next day, they were washed with PBS and unmasked in 1:100 diluted antigen retrieval solution (Vector). Embryos were then washed again in PBS for 10 min (RT) followed 30 min in Dent fixative (20% DMSO/methanol) at 4°C. Subsequently, all samples were bleached overnight at 4°C in 5% H_2_0_2_/20% DMSO/methanol. Next, embryos were blocked overnight in 10% FBS/20%DMSO/PBS at 4°C and then incubated in anti-neurofilaments antibody clone 2H3 for 4 days (dilution 1:100 in blocking solution) followed by secondary HRP-conjugated antibody (1:1000) for 24 hours. Then, embryos were washed in 20% DMSO/PBS and incubated in 0.6% tween/PBS overnight. Finally, samples were washed in PBS and incubated in 0.05% DAB for 2 hours. H_2_0_2_ was then used as a substrate. Labeled embryos were washed in PBS and cleared in ascending glycerol concentration (20%, 40%, 60%, 80%). They were stored in 80% glycerol at 4°C. Images were captured using Nikon SMZ18 stereomicroscope and post-processed in Helicon focus to create sharp images. Surface area occupied by a given nerve was measured in ImageJ. Axons were traced in Neurolucida 360.

### Diolistics and DiI tracing

We used DiOlistic approach using Gene Gun helium-powered system from Biorad. Bullets were prepared as described (Sherazee and Alvarez, 2013), tubing was coated with 10 mg/ml polyvinylpyrollidone (PVP). We mixed 100 mg Tungsten beads and 2.5 mg DiI or DiO (dissolved in CH_2_Cl_2_). After CH_2_Cl_2_ evaporation, resulted powder was transferred into aluminum-wrapped falcon tube and 3 ml H_2_0 were added. Solution was sonicated at 4°C until no clumps were visible (30 – 45 minutes). Then, solution was sucked into tubing in prep station, beads were able to settle down and water was removed. Tubing was rotated 1 hour during continuous drying with nitrogen (2-3 liters/min). Finally, 1.3 cm bullets were cut from tubing and stored in 4°C with silicagel beads to prevent rehydration.

Slices for DiOlistics were prepared as follows: mice were perfused with 20 ml 4% PFA/PBS, brains were isolated and postfixed 30 minutes in 4% PFA/PBS. Then, brains were washed 1 – 3 hours in PBS and 20 minutes in 15% sucrose/PBS following another 20 minutes in 30% sucrose/PBS. 250 μm coronal slices were prepared using vibratome Leica 2000S. Prior to shooting, slices were treated 5 minutes in 15% sucrose and 5 minutes in 30% sucrose. Dye was carried using pressure 120 Psi and modified filter as described (Sherazee and Alvarez, 2013). After shooting, slices were washed 3x in PBS quickly and dye was let to diffuse 40 min at 4°C. Slices were then mounted onto glass slice in 0.5% n-propylgallate/90% glycerol/PBS (NPG) and imaged by CARV II/Nikon Ti-E spinning disc. For carbocyanine dye tracing mice were perfused transcardially with PBS and fixed with 4% PFA/PBS. Next, either small DiI crystal was placed or 0.1 μl of DiI solution was injected into target area. We used 2.5 mg/100 μl concentration, DiI was dissolved in DMSO. Slices were prepared in vibratome and scanned by Leica TCS SP8. Details are as follows.

Corpus callosum tracing: DiI solution was injected into superficial layers of cortex. Brains were maintained in 4%PFA/PBS in 37°C for 3 – 4 weeks. Then, brains were cut in oblique (horizontal+20°) direction (Fig. S3a), 150 µm-thick sections were prepared. Slices with traced axons were mounted onto glass slide in NPG mounting solution.

Hippocampo-septal axon tracing: After fixation, brains were trimmed to expose septum. DiI crystals were inserted into the medial septum and brains were maintained in 4%PFA/PBS for 1-2 weeks. 100 µm (P0-1) or 150 µm (P8) coronal slices were prepared to screen for retrogradely labelled CA1 neurons in the hippocampus.

Visual corticospinal axon tracing: DiI solution was injected into primary visual area. After 2 – 3 weeks, brains were cut sagitally to 150 – 180 µm slices, that were mounted in NPG with Hoechst. The tracing pattern was compared with data from Allen brain atlas conectivity studies to ensure that we targeted correct area. We observed two axon branches in the diencephalon: first branch growing into pyramidal tract (corticospinal axons) and second to superior colliculus (collicular axons). To quantify the axonal growth into the pyramidal tract, we compared fluorescence intensity of corticospinal axons vs. intensity of axons before branching. We refer to this ratio as visual pruning index (VP index), with its lower values indicating presence of refinement.

### In utero electroporations

*In utero* electroporations were done as previously described (Haddad-Tovolli et al., 2013). Briefly, pregnant mice were anaesthetized by 2,5% isoflurane. Anaesthesia was maintained by 2% isoflurane. We injected pCAGGS-EGFP plasmid (3 μg/μl) into ventricles of E14.5 embryos (migration assay, analysis at E17.5) or E15.5 embryos (callosal axons, analyzed at P6). Electroporation was carried out by small paddle electrodes (35 V, 5 pulses, 950 msec interval) to target sensory cortex. For migration assays, embryos were harvested at E17.5, brains were fixed in 4% PFA/PBS, sliced (150 μm vibratome sections), counterstained with Hoechst and scanned by Leica TCS SP8. In this case only, both WTs and *crmp2^+/-^* embryos were used as controls. Callosal axons were analyzed at P6. After birth, pups were nurtured by a foster mother. At P6, mice were sacrificed, brains were fixed in 4% PFA/PBS, sliced (150 μm vibratome sections) and analyzed by CARV II/Nikon Ti-E spinning disc.

### Semaphorin assays and live imaging

Mouse E16.5 hippocampal neurons were prepared and cultured as described (Balastik et al., 2015). Briefly, pregnant mice were sacrificed, embryos isolated and decapitated in cold HBSS with 10 mM Hepes. Hippocampi were isolated, moved to Neurobasal medium (Neurobasal (Gibco) with 2,5% Β27 supplement (Gibco), 2.5 mM glutamine and 1% Penicillin/Streptomycin solution) and triturated. Subsequently, solution was strained through a 40-μm strainer (Biologix) and spun down (300g, 2 min., 4°C). Supernatant was removed, remaining cells resuspended in a fresh Neurobasal medium and counted. Neurons were plated into 24-glass bottom plates (100 000 cells/well) coated with laminin (1 μg/ml) and poly-D-lysine (50 μg/ml) and cultured in Neurobasal medium that was refreshed every two days. Neurons were transfected with pCAGGS-EGFP using lipofectamine-2000 (Invitrogen), 24 hours after plating. Mouse Semaphorin 3A and 3F (carrier free) or control Fc were purchased from R&D systems and were diluted to 1 mg/ml stock concentration. At DIV 7, medium volume was adjusted to 300 μl in each well. Neurons were photographed three times (with 20 minutes gaps), then 50 μl of fresh medium with Fc or Sema3A (final concentration 1 nM) or Sema3F (final concentration 5 nM) was added. Subsequently, neurons were imaged every 20 minutes for 16 hours by Leica DMI6000 equipped with a heating box and CO_2_ atmosphere.

### Microfluidic chambers

Microfluidic chambers were prepared as described before (Ionescu et al., 2016, Maimon et al., 2018). E11.5 – E12.5 spinal cord explants were dissected in cold HBSS and placed into Laminin (3 μg/ml) and Poly-L-Ornithine (1,5 μg/ml) coated proximal well. After 3 – 4 days, axons entered the distal compartment. Then, explants were labelled by Alexa 647-conjugated choleratoxin subunit B. 5 nM Sema3A of control Fc was applied distally and axons were photographed by Leica DMI6000 microscope every 10 minutes during 14 hours interval. Axons growing at least 50 μm were analyzed.

### Analysis of developmental motor axon pruning

WT and *crmp2^−/−^* pups (both sexes) were sacrificed on P11, the thorax was excised as previously described (Kleele et al., 2014, Brill et al., 2013, Marinkovic et al., 2015, Kerschensteiner et al., 2008) and fixed in 4 % PFA in 0.1 M phosphate buffer (PB) for 1 hour on ice. The triangularis sterni muscle was dissected and incubated overnight (4°C) in anti-βIII-tubulin antibody conjugated to Alexa488 (BioLegend 801203; 1:200) and Alexa594-conjugated α-bungarotoxin (Invitrogen B13423; 50 µg/ml; 1:50) in blocking solution (5 % BSA, 0.5 % Triton X-100 in 0.1 M PB). Muscles were then washed in 0.1 M PB and mounted in Vectashield (Vector Laboratories). Z-stacks were recorded using a confocal microscope (FV1000, Olympus) equipped with a 20x/0.8 N.A. oil-immersion objective and analyzed for the percentage of doubly innervated NMJs using Fiji (cell counter plugin) (Schindelin et al., 2012).

### Behavioural tests

1. Ultrasonic vocalizations were recorded from P6, P8 and P12 pups (WT n=14, *crmp2^−/−^* n=13). Each pup was taken from its home cage, put into Styrofoam box and recorded for 5 minutes with microphone (Dodotronic Ultramic 250K, Italy) placed at the top of the box. Audacity software (freely available) was used for recordings with sampling frequency set to 250 kHz. The vocalizations were analyzed automatically using Avisoft-SASLab Pro, however, the automatic analysis was checked and manually corrected if necessary. Main parameters measured were number of vocalizations and its total length.
2. The Elevated plus maze (EPM) is a cross-shaped maze elevated 40 cm above the floor level, with all four arms (30 cm long, 5 cm wide, 16 cm high) accessible from the central platform. Two opposite arms were enclosed by opaque walls (closed arms), while the other two were without walls (open arms). The test assesses spontaneous and anxiety-like behaviors of the animals. The closed arms are perceived as safer by the animals, as they are darker, more protected and without risk of falling. Anxious animals are expected to spend most of the time in the closed arms, while less anxious and more explorative individuals explore the open arms more often. The behavior of the animals during each 5-min session was recorded by a web camera placed above the apparatus. Locomotor activity (total distance) and time spent in different compartments was analyzed offline by digital tracking software (Ethovision, Noldus). Numbers of animals: n=10 for both WT and knockouts, aged 31 – 44 weeks.
3. Spontaneous alternation was tested in three-armed maze (Y-maze), with each arm 35 cm long, 6 cm wide and 18 cm high. The mice were left free to explore the empty apparatus for 8 min. Between trials, the apparatus was cleaned by ethanol and then wiped clean and dry to erase any scent marks. Number of arm visits was counted, indicating the exploratory activity (a visit was counted if the mouse placed all paws into the arm). Spontaneous alternation was measured as the ratio of actual triads (three different arms entered in three subsequent visits) to potential triads (theoretical maximum performance). In the Y-maze task, 17 male mice (9 wt, 8 ko), aged 31 – 44 weeks, were tested.
4. The active place avoidance task, or AAPA (Bures et al., 1997, Burghardt et al., 2012) (for review see (Stuchlik et al., 2013)) is a rodent cognitive task testing hippocampal functions, including spatial navigation, cognitive coordination and flexibility. As a dry-arena task, it is more suitable for mice than the Morris water maze, as mice are worse swimmers than rats and tend to be overtly stressed by water immersion (D’Hooge and De Deyn, 2001). We used a carousel maze (circular arena 56 cm in diameter) with electrified grid floor, surrounded by a transparent plexiglass wall, rotating at approximately 1 rotation per minute. The apparatus was located in a dimly lit room with abundant extramaze cues, with and additional, highly contrast cue card in close proximity (1.5 m) of the arena. A computer-based tracking system (Tracker, Biosignal Group, USA) recorded the positions of the mouse and the arena at a sampling rate of 25 Hz. A 60-deg unmarked to-be-avoided sector was defined in the coordinate frame of the room by tracking software. Each entrance into the sector was punished by mild electric foot shocks (scrambled; 100 HZ alternating current; 40 – 80 V) delivered by the tracking system into the grid floor. Each shock lasted 0.5 s, and was repeated after 0.9 s if the mouse failed to escape the to-be-avoided sector in time. Intensity of the shock was individualized for each mouse (0.2-0.4 mA), to ensure escape reaction while avoiding excessive pain. The training schedule consisted of 5 acquisition sessions and 4 reversal sessions, where the sector position was changed by 180 degrees. Two 10-min sessions were scheduled for each experimental day, separated by approximately 3 hours of rest in the home cage. The arena rotated while the sector remained fixed in the reference frame of the room, therefore, the mice had to move actively away from the sector in the direction opposite to arena rotation, otherwise they would be carried into the sector. For successful avoidance, the animal had to separate the distant room-frame cues, which could be used to locate and avoid the sector, from the irrelevant, arena-frame cues. Selection of the correct spatial cues and achievement of the correct behavioral strategy requires segregation of spatial frames, a skill that is considered equivalent of human cognitive coordination (Wesierska et al., 2005). Furthermore, cognitive flexibility was tested in reversal sessions, to adjust the behavioral response to reversed sector location.

The trajectory of the mice was analyzed offline using the custom-made and freely-available Carousel Maze Manager 4.0 (Bahník, 2014). Total distance walked, a measure of locomotor activity of the mouse, and the number of entrances, a measure of its ability to avoid entering the to-be-avoided sector, were used as the most important output parameters. Active place avoidance performance was evaluated by two consecutive RM-ANOVA analyses, with sessions taken as repeated measures, and genotype as a between-subject factor. In this task, 21 male mice (11 ko, 10 wt), 12 - 14 weeks old, were tested.

### Light Microscopy

5 types of microscopes were used in this study. 1) Leica TCS SP8 confocal equipped with 405 nm laser (Hoechst), 488 nm laser (DiO, EGFP excitation), 552 nm laser (DiI excitation), 10x/0.3 dry and and 25x/0.75 immerse-oil objectives (Fig. 2g, Fig. 4, Fig. 6d, Fig. S1f, Fig. S3c, Fig. S4b). 2) Inverted fluorescent microscope Leica DMI6000 equipped with 20x/0.4 dry objective, 37°C incubator and CO_2_ chamber for live imaging (Fig. 1f, Fig. 5). 3) CARV II/Nikon Ti-E spinning disc equipped with 20x/0.5 dry, 40x/1.3 and 100x/1.4 immerse-oil objectives (Fig. 1b, Fig. 2c and d, Fig. 3, Fig. 6a, b, f, Fig. S1a, f, Fig. S4a, Fig. S5, Fig. S6a). 4) Olympus FV1000 confocal microscope equipped with 20x/0.8 objective (Fig. S4c). 5) Nikon SMZ18 stereomicroscope for microdissections and whole-mount analysis equipped with 1x and 2x SHR Plan Apo objectives (Fig. 1c, Fig. 2a, Fig. S1c and d, Fig. S2). Microscopy images from figures 1, 2, 3, and 4 and figures 1, 3 and 4 are compositions of more fields automatically generated by Leica LAS X software or NIS elements software. Maximum projections were generated in ImageJ.

### Image processing and statistics

We used Neurolucida 360 software to reconstruct callosal axons from both *in utero* electroporation and DiI-tracing experiments. In case of oblique sections, we picked only midline regions (approximatelly 100 μm from midline to each side) and traced axons semi-automatically using AutoNeuron algorhytm. To display overall axon growth direction, polar histograms and fan in diagrams were generated using Neurolucida explorer. Polar histograms show total axon length in a specific degree range. We counted lengths in each 20° of histogram. Fan in diagrams represents the same in another graphical view when all traced axons rise from a single point. Dendrites and dendritic spines were analyzed either with Neurolucida 360 or with NeuronJ plugin in ImageJ. Images from whole-mount preparations were processed in Helicon focus to create sharp projections. Growth of peripheral nerves and all data from time-lapse imaging series were analyzed in ImageJ. Neuron migration was analyzed in Image J using cell counter plugin. Measurements of axon pruning from immunohistochemistry and DiI-tracing experiments were done in ImageJ. In all figures, different channels of image series were combined in pseudo-colour using the „screen” function in Adobe Photoshop, and adjusted to enhance low-intensity objects. For comparison of two independent groups, unpaired t-test was used unless stated otherwise. Data from behavioural experiments were analyzed by unpaired t-test (P6, P8 vocalization, Y maze, EPM), ANOVA (AAPA) or Kruskal-Wallis test (P12 vocalization). All data are presented as means±SD, unless stated otherwise.

## ACKNOWLEDGEMENTS

The work of MB, JZ, MJ, TP, BP and AS was supported by Czech Health Research Council grant no. NV18-04-00085. MB, RW, BP, JZ, MK were supported by Czech Science Foundation grant no. 16-15915S. The work of JZ, RW and BP was supported by Grant Agency of the Charles University grants no. 682217, 524218 and 1062216 respectively. MJ, TP and AS were supported by Czech Science Foundation grant no. 19-03016S and Czech Health Research Council grant no 17-30833A. MW, MSB and TM was supported by the ERC (FP/2007-2013; ERC Grant Agreement n. 616791), the German-Israeli Foundation, SFB 870 and the Munich Cluster for Systems Neurology (SyNergy; EXC 1010). RS was supported by RVO 68378050 by Academy of Sciences of the Czech Republic and by LM2015040 (Czech Centre for Phenogenomics), CZ.1.05/2.1.00/19.0395, CZ.1.05/1.1.00/02.0109 funded by the Ministry of Education, Youth and Sports and the European Regional Development Fund. We acknowledge the Microscopy Centre - Light Microscopy Core Facility, IMG ASCR, Prague, Czech Republic, supported by MEYS (LM2015062), OPPK (CZ.2.16/3.1.00/21547) and (LO1419), and Light Microscopy Core Facility, IPHYS ASCR, Prague, Czech Republic, for their support with confocal and live imaging.

## AUTHOR CONTRIBUTION

Study design: JZ, MB; data acquisition and analysis: JZ, RW, KJ, MJ, RM, TP, BP, MW, MSB, MK, PK, XZ and MB; supervision: GAB, RS, TM, AS, EP, MB; writing - original draft: JZ, MB; writing – review and editing: JZ, GAB, RM, AS, TP, MSB, TM, MB; project administration: MB

## DECLARATION OF INTEREST

The authors declare no competing interest.

## SUPPLEMENTAL INFORMATION

**Table S1.** Number of analyzed animals/replicates, related to methods.

| analysis | Figure | WT | KO | notes |
| --- | --- | --- | --- | --- |
| E10.5 - 11.5 n. V | **1C, D** | 10 | 10 | embryos from the same litter were compared |
| E10.5 - 11.5 spinal nerves | **1C, E** | 8 | 7 |  |
| E12.5 n. V | **S2B, C** | 4 | 5 |  |
| E17.5 IUE migration | **S1F** | 6 (wt 2, het 4) | 4 |  |
| ventricles | **S1D, E** | 3 | 3 |  |
| IUE corpus callosum | **2D, F** | 6 | 3 |  |
| Dil tracing callosum P9 | **2G, I** | 6 | 9 |  |
| Dil tracing callosum adults | **S3C, D** | 9 | 7 |  |
| callosum P30 | **2B** | 3 | 3 |  |
| callosum adults | **2A, B** | 6 | 6 | coronal/sagittal: WT 3/3 vs. KO 3/3 |
| IPB pruning | **3G, S4A** | 6 | 7 | coronal/sagittal: WT 4/2 vs. KO 5/2 |
| visual CS axons P9 | **4A, C** | 3 | 3 |  |
| visual CS axons adults | **4A, C** | 5 | 5 |  |
| Dil tracing H-septal P1 | **S4B** | 3 | 3 |  |
| Dil tracing H-septal P8 | **S4B** | 3 | 3 |  |
| NMJs P11 | **S4C, D** | 3 | 3 |  |
| DG spines P30 | **6A, C** | 3/61 dendrites | 3/64 dendrites |  |
| DG spines adults | **6B, C** | 3/41 dendrites | 3/37 dendrites |  |
| DG dendrite branching | **6D, E** | 3/27 dendrites | 3/25 dendrites |  |
| PFC spines juvenile | **6F, G** | 3/51 dendrites | 3/54 dendrites |  |
| PFC spines adults | **6F, G** | 3/53 dendrites | 3/57 dendrites |  |
| CA1 spines adults | **S6A, B** | 3/36 dendrites | 3/36 dendrites |  |
| behavior - vocalization | **7A, B, C** | 14 | 13 |  |
| behavior - Y maze | **7D, E** | 9 | 8 |  |
| behavior - EPM | **7F, G, H** | 10 | 10 |  |
| behavior - AAPA | **S6C** | 10 | 11 |  |
| in vitro - MFC, E12.5 explants | **1F, G** | 3 experiments/7 control vs 8 Sem3A explants | 3 experiments/8 control vs 8 Sem3A explants |  |
| in vitro - retraction assay | **5A, C** | 3 experiments/4 control vs 7 Sem3A vs 6 Sem3F wells | 3 experiments/5 control vs 6 Sem3A vs 5 Sem3F wells |  |
| in vitro - WB | **5D, E** | 3 experiments/triplicates |  |  |

**Figure S1.**
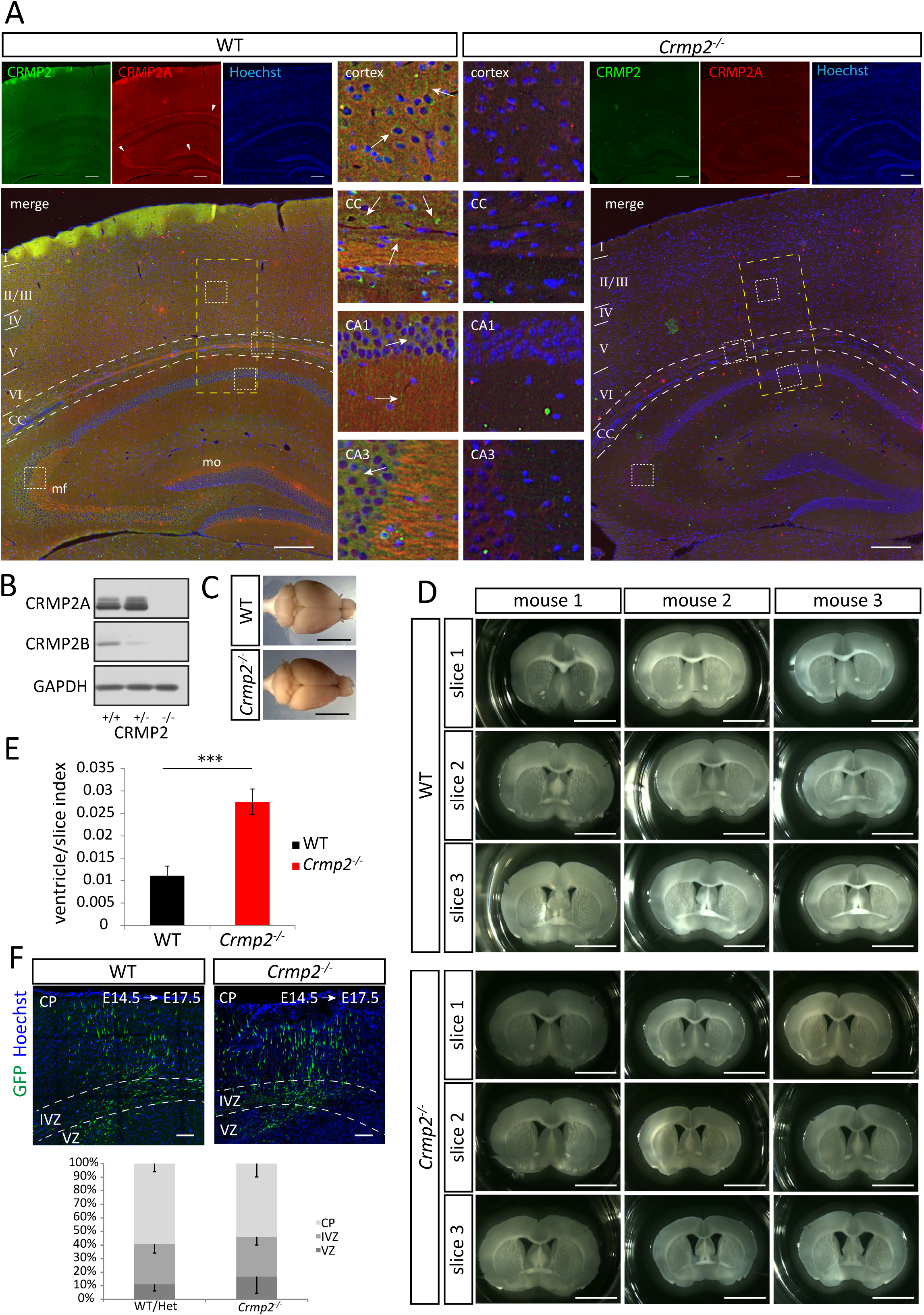
Related to Figure 1. Analysis of the brain morphology of *crmp2^−/−^* mice. (A) CRMP2 (A+B isoforms, arrows) and CRMP2A (arrowhead) isoform immunostaining of coronal brain sections from adult WT and *crmp2^−/^*^−^ mice. Both isoforms are missing in *crmp2^−/^*^−^ mice. Yellow rectangles depict the area shown in Figure 1B. CRMP2B is present throughout the cortex, and hippocampal CA1 and CA3 regions. CRMP2A is localized mostly in callosal axons (CC), mossy fibers (mf) and inner molecular layer (mo) of dentate gyrus. (B) WB analysis of brain lysates of *crmp2* ^+/+^, ^+/-^ and ^−/−^ mice. (C) Upper view of WT and *crmp2^−/^*^−^ brains showing similar brain size. (D) Three 250-μm coronal sections from 3 WT and *crmp2^−/−^* mice demonstrate enlarged ventricles in *crmp2^−/−^* mice. (E) Quantification of the ventricle areas normalized to total slice area (ventricle/slice index, WT 0.011±0.001, *crmp2^−/−^* 0.027±0.003, p<0.001), means±SD are shown. (F) Analysis of cortical neuron migration (WT/*crmp2^+/-^* n=6, *crmp2*^−/−^ n=4). Both WTs and heterozygous (Het) embryos were used as controls. Embryos were electroporated with GFP expressing plasmid at E14.5 and analyzed at E17.5. Maximum projections are shown. Quantification of percentages of GFP positive neurons in different layers is shown. CP – cortical plate, IVZ – interventricular zone, VZ – ventricular zone. ***p<0.001, t-test. Scale bars: 100 μm in A and F, 5 mm in C and D.

**Figure S2.**
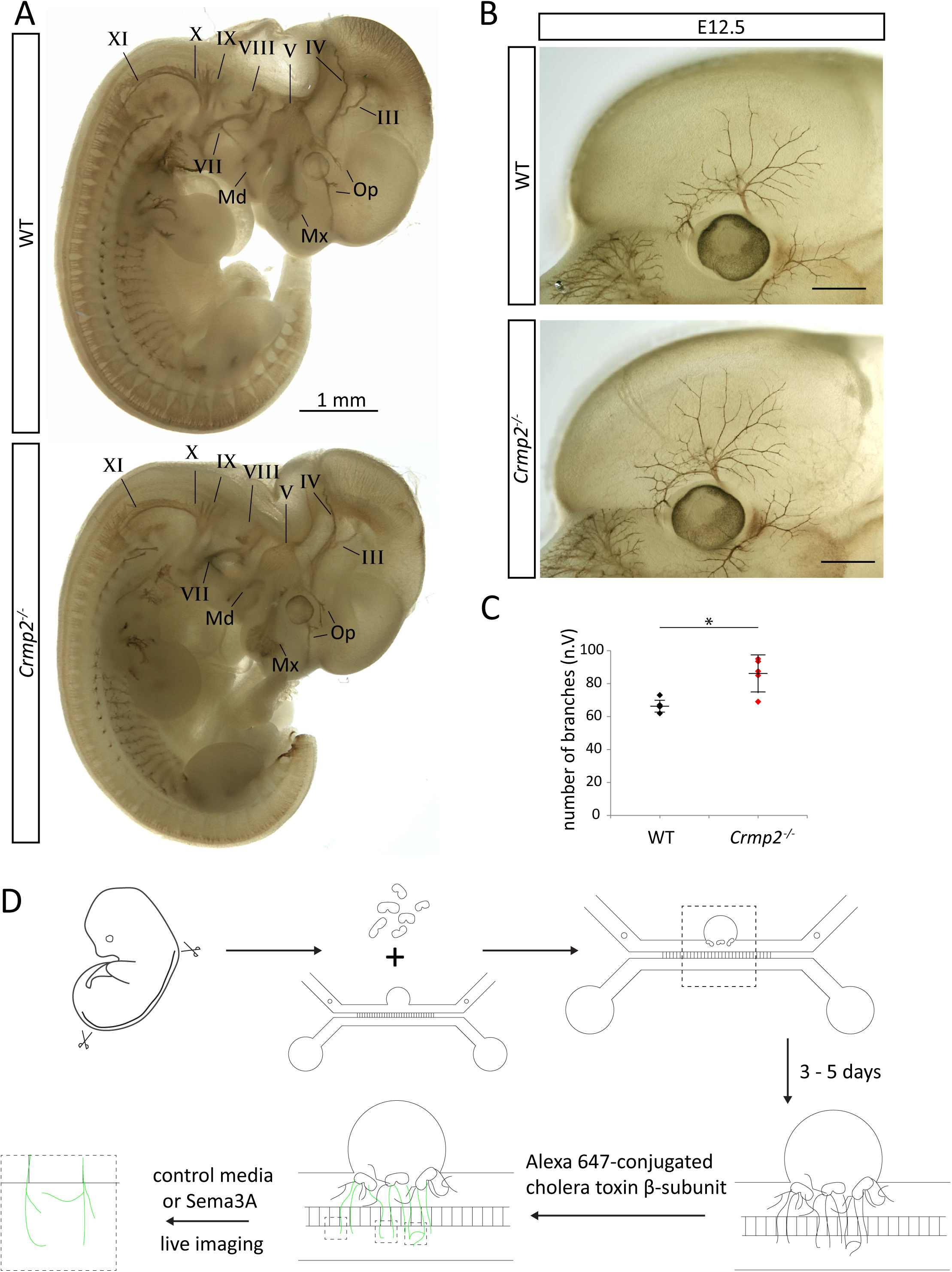
Related to Figure 1. Peripheral nerve growth and motor neuron cultures in microfluidic chambers. (A) WT and *crmp2^−/−^* E11.5 embryos stained with antibody against neurofilaments. All peripheral nerves are present in *crmp2^−/−^*. III – occulomotor, IV – trochlear, V – trigeminal, VII – facial, VIII – vestibulocochlear, IX – glossoparyngeal. X – vagal, XI – accessory nerve. Op – ophthalmic branch, Mx – maxillar branch, Md – mandibular branch of the trigeminal nerve (B) Whole-mount immunolabeling of embryos at E12.5 (WT n=4, *crmp2^−/−^* n=5, 2 litters) note the increased branching of the Op branch in *crmp2^−/−^*. (C) Quantification of the number of branches (WT 67±4.5, *crmp2^−/−^* 86±10.3, p<0.05), means±SD are shown. (D) Experimental design of motor neuron cultures in microfluidic chambers using spinal cord explants. Dashed squares in the second row show areas that were scanned and analyzed. * p<0.05, t-test. Scale bars: 1mm in A, 500 μm in B

**Figure S3.**
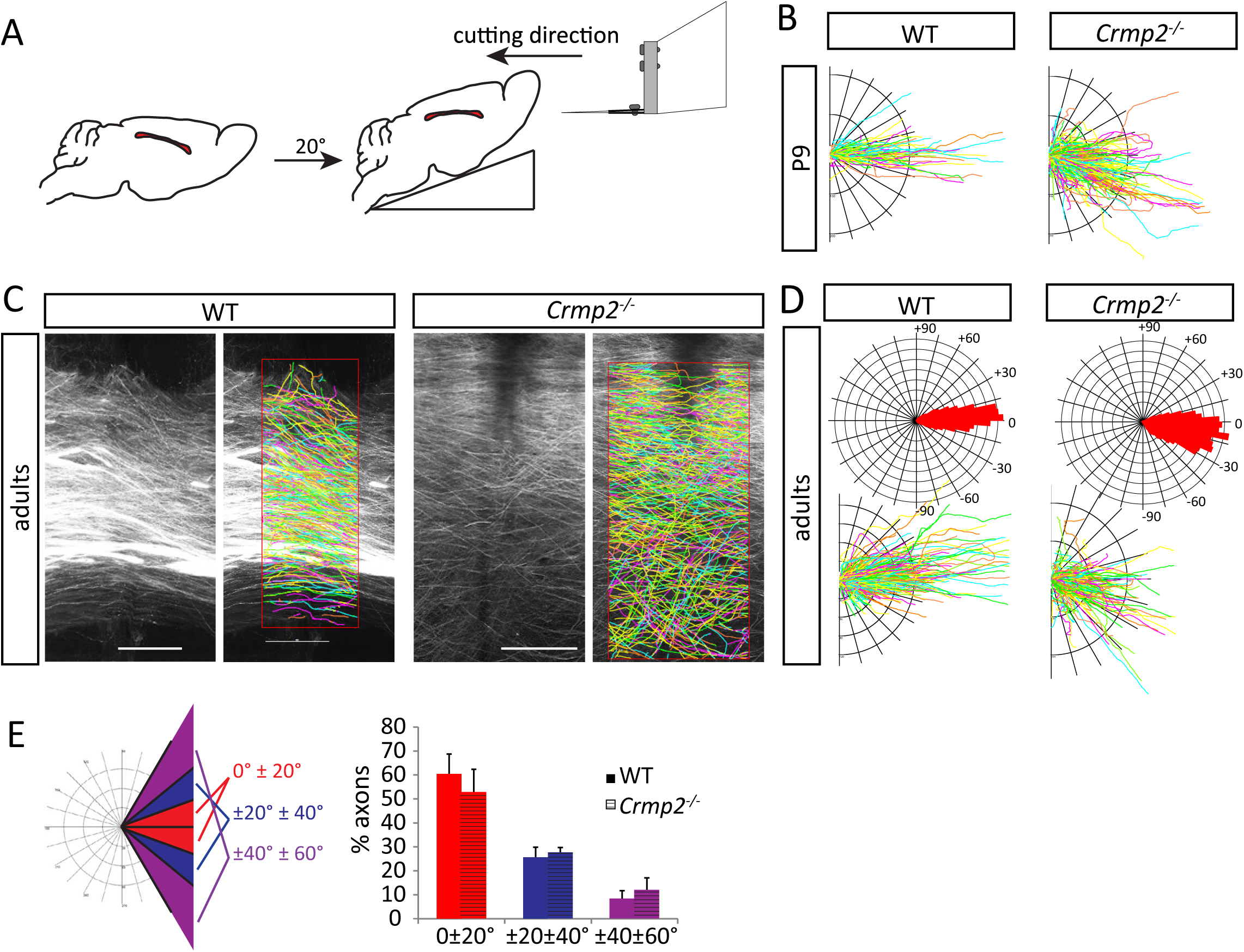
Related to Figure 2. Analysis of the guidance of callosal axons in oblique sec ons. (A) Drawing of the experimental design used to prepare large ons of corpus callosum in one slice. (B) Fan in diagrams from tracing of P9 callosal axons. (C) DiI tracing of callosal axons from oblique ons in the midline of the adult WT (n=9) and *crmp2^−/−^* (n=7) mice reconstructed in Neurolucida360. (D) polar histograms and fan in diagrams from reconstructed adult callosal axons. (E) Quantification of axon orienta n in the midline of corpus callosum revealed no differences between WT and knockout mice (WT, 0°-±20°: 60.5±8.23%; ±20°-±40°: 25.6±4.18%; ±40°-±60°: 8.24±3.47%; *crmp2^−/−^*, 52.7±10%, p=0.1, 27.5±2.23%, p=0,3, 12.1±5%, p=0.09). Means±SD are shown, scale bars: 200 μm.

**Figure S4.**
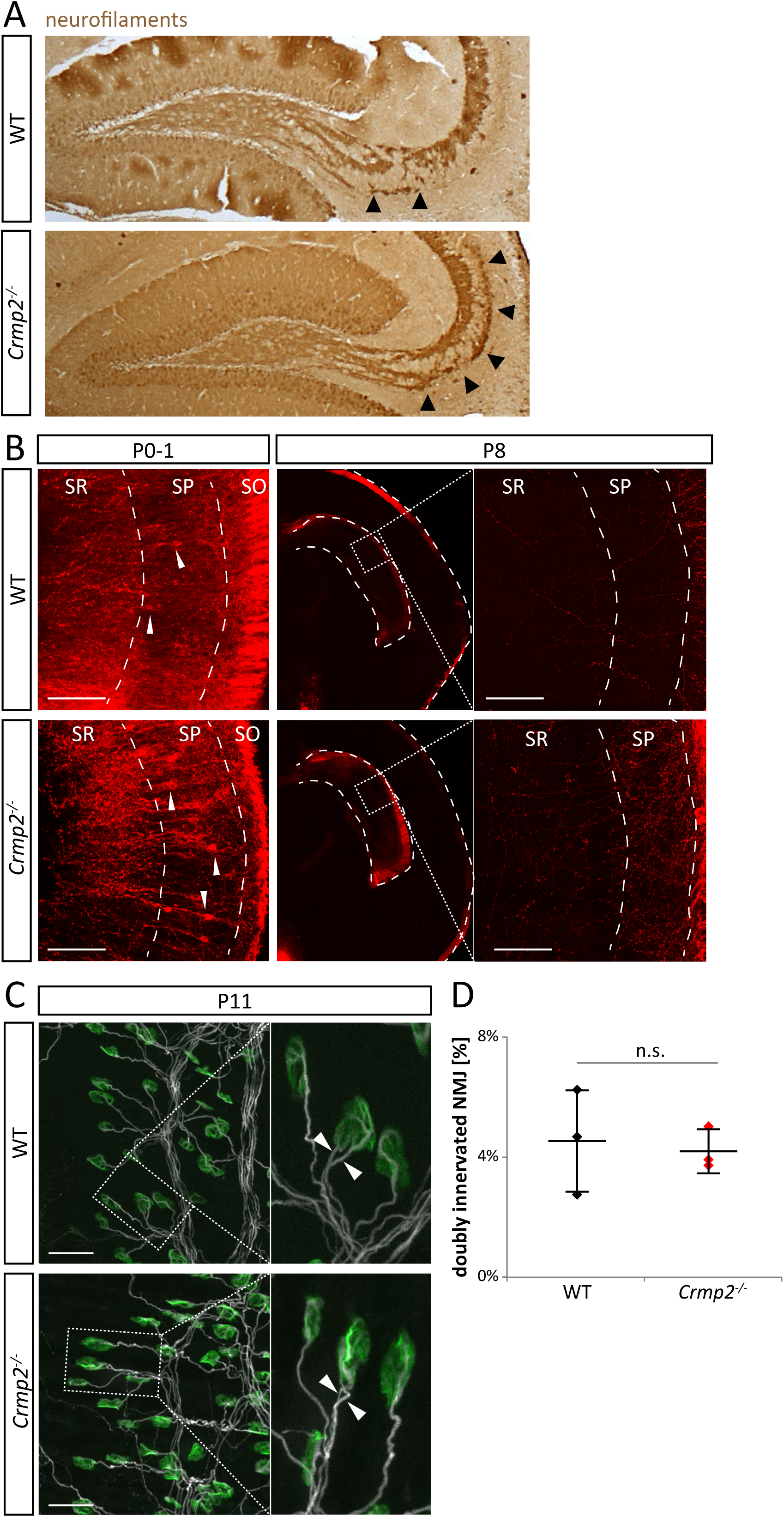
Related to Figure 3. Axon pruning in *crmp2^−/−^* mice is altered in Sema3F-related but not Sema3A-related regions. (A) Calbindin immunostaining of sagittal brain sections show defects in pruning similar to those seen in coronal sections of WT and *crmp2^−/^*^−^ mice in Fig. 4. (B) Hippocamposeptal axons are pruned in both WT and *crmp2^−/^*^−^ mice (n=3). At P0-1, regrogradely DiI-labeled CA1 neurons showed strong signal in both WTs and *crmp2^−/−^* indicating presence of CA1 projections into the medial septum. Arrowheads in the left column show retrogradely labeled cell bodies. Conversely, at P8, no retrogradely labeled cell bodies were detected in CA1 granular layer in either WT or *crmp2^−/−^* mice. Squares in the second column are shown in a higher magnification in the third column. SR – stratum radiatum, SP – stratum pyramidale, SO – stratum oriens (C) Maximum intensity projections of P11 triangularis sterni muscle NMJs of WT and *crmp2^−/−^* mice (n=3). Axons were stained with βIII-tubulin antibody (white), postsynaptic acetyl choline receptors with α-bungarotoxin (green). Boxed areas are shown on the right at higher magnification with examples of doubly-innervated NMJs where two axons are competing for synaptic innervation (arrowheads). (D) Quantification of the number of doubly-innervated NMJs revealed no significant difference (WT 4.5±1.8%, *cmrp2^−/−^* 4.2±0.7%, p>0.99). Mann-Whitney-test, means±SD are shown, scale bars: 100 μm in B, 50 μm in C.

**Figure S5.**
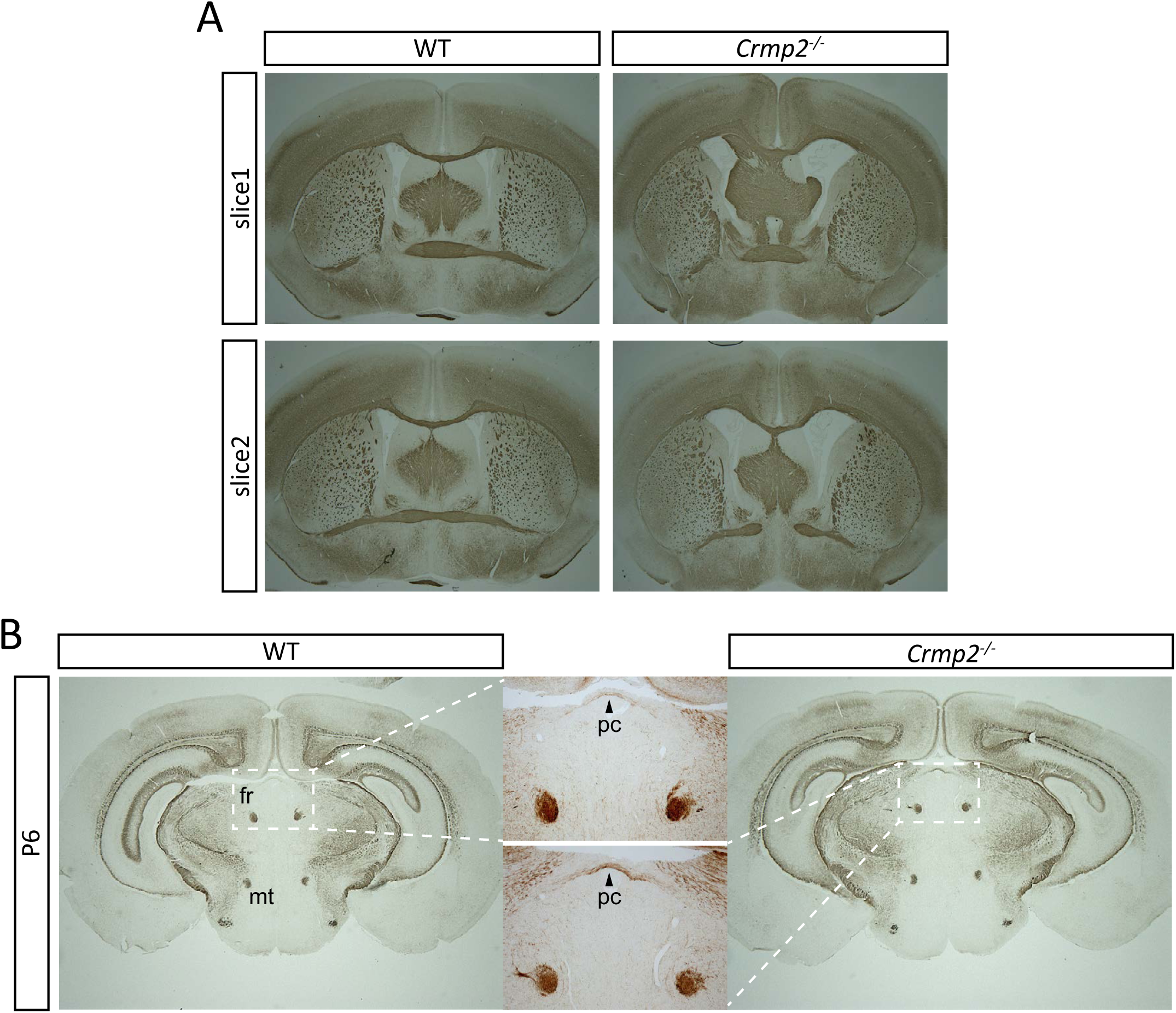
Related to Figure 4. Growth of anterior commissure and retroflex fascicle (Meynert’s) is unaffected in *crmp2^−/−^* mice. (A) At P14, anterior commissure is present in both WT and *crmp2^−/−^* mice. Two consecu coronal sec ons labeled with an body against neurofilaments are shown. (B) Retroflex fascicle (fr), as well as mammilothalamic tract (mt) are normally formed in *crmp2^−/−^* mice. Coronal sec ns at the level of posterior commissure (pc) are shown.

**Figure S6.**
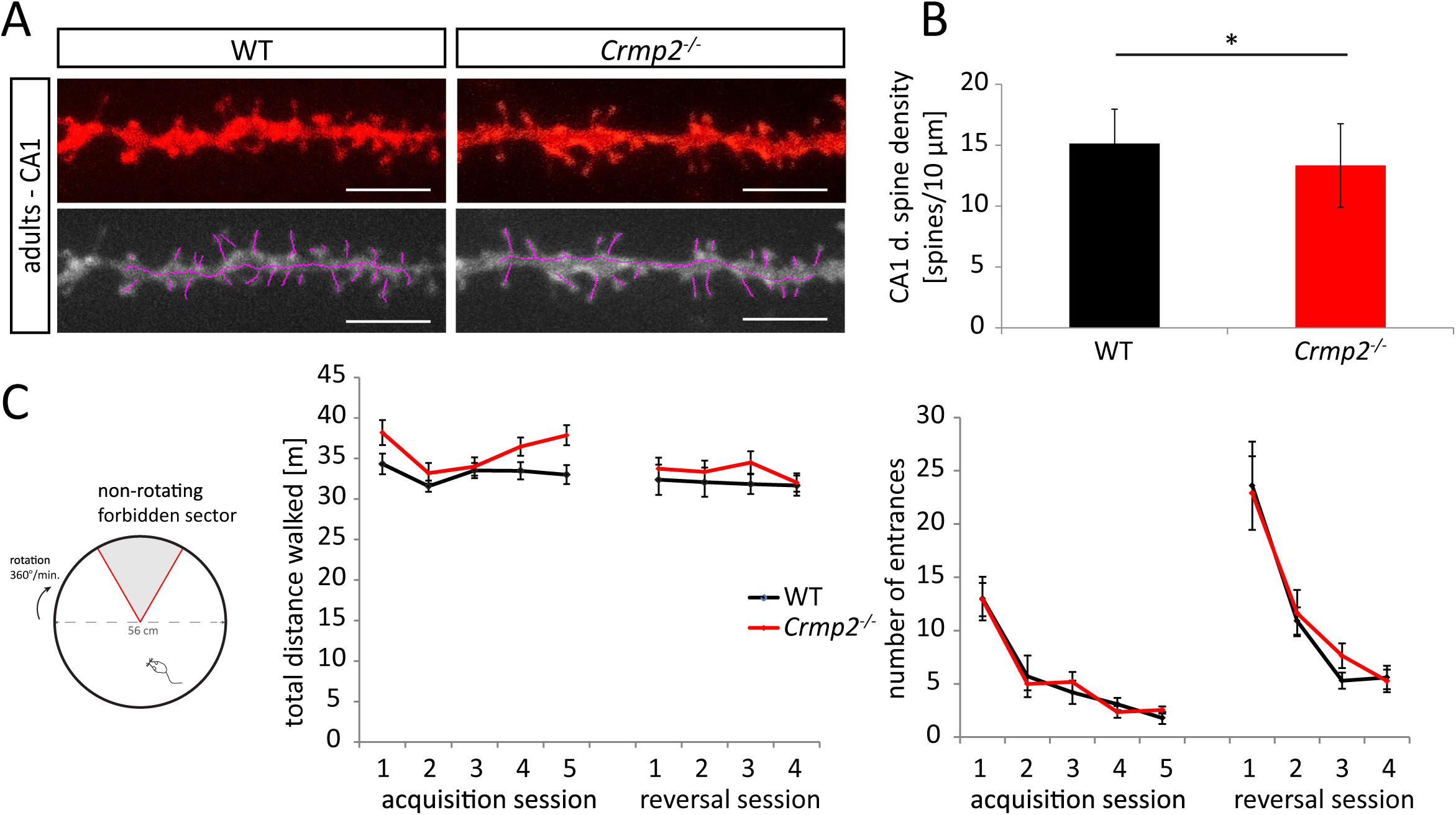
Related to Figures 6 and 7. Dendri c spine density in hippocampal CA1 region and behavioral analysis of hippocampus-related spa al cogni ve abili es using AAPA task. (A) Analysis of DiOlis cally labeled dendri c spine density in hippocampal CA1 region, secondary apical dendrites (n=3, 36 dendrites). (B) Quantification of the spine density in CA1 neurons in (A). The density is decreased in *crmp2^−/−^* mice (WT 15.2±2.8 spines/10 μm, crmp2^−/−^ 13.3±3.4 spines/10 μm, p=0.02). (C) AAPA (Ac Allothe c Place Avoidance) task for spa l cogni abili es. WT as well as knockouts learned the task successfully as demonstrated by a decreasing number of entrances into the sector during both acquisi on and reversal phase. No significant difference was measured between WT and *crmp2^−/−^* mice in either acquisi on or reversal setup (see material and methods) *p<0.05, t-test. Scale bars: 5 μm.

